# Dual-use virulence factors of the opportunistic pathogen *Chromobacterium haemolyticum* mediate haemolysis and colonization

**DOI:** 10.1101/2025.01.10.632398

**Authors:** Leo Dumjahn, Philipp Wein, Evelyn M. Molloy, Kirstin Scherlach, Felix Trottmann, Philippe Meisinger, Louise M. Judd, Sacha J. Pidot, Timothy P. Stinear, Ingrid Richter, Christian Hertweck

## Abstract

*Chromobacterium haemolyticum* is an environmental bacterium that can cause severe and fatal opportunistic infections in humans and animals. Although *C. haemolyticum* is characterized by its strong β-haemolytic activity, the molecular basis of this phenotype has remained elusive over the more than fifteen years since the species was first described. Here, we report the discovery of a family of cyclic lipodepsipeptides that are responsible for the potent haemolytic activity of *C. haemolyticum.* Comparative genomics of *Chromobacterium* spp. isolated from different environments revealed a completely conserved gene locus (*chl*) encoding a non-ribosomal peptide synthetase (NRPS). Metabolic profiling of *C. haemolyticum* DSM 19808 identified a suite of cyclic lipodepsipeptides as the products of the *chl* locus, with the three main congeners (jagaricin, chromolysin A and B) being elucidated by a combination of tandem mass spectrometry, chemical derivatization, and NMR spectroscopy. Significantly, a *C. haemolyticum chl* deletion mutant is devoid of haemolytic activity. Moreover, purified jagaricin, chromolysin A and B are haemolytic at low-micromolar concentrations in an erythrocyte lysis assay. Further bioassays demonstrated that the cyclic lipodepsipeptides are crucial for biofilm-forming and swarming behavior of *C. haemolyticum*. MALDI mass spectrometry imaging showed that primarily chromolysin A and B are involved in these processes *in vitro*. Our data shed light on the bioactivities of chromolysin A and B, specialized metabolites that likely contribute to both successful colonization of new niches and virulence potential of *C. haemolyticum*.

**Importance:** Despite the rising incidence of *Chromobacterium haemolyticum* as a serious opportunistic pathogen, there is limited information on whether the competitive traits that ensure its survival in its freshwater niche contribute to host infection. We present a case where bacterially produced specialized metabolites act as lynchpin chemical mediators that are not only responsible for the pronounced haemolytic phenotype of *C. haemolyticum* but are crucial for biofilm formation and swarming motility. These results exemplify a case of coincidental evolution, wherein the selective pressures encountered in a primary environmental niche drive the evolution of a trait impacting virulence. This knowledge provides a foundation for the development of antivirulence therapies against the emerging pathogen *C. haemolyticum*.

## Introduction

Environmental microbes interact with a broad range of microorganisms through cooperation and competition (1, 2). Survival likely requires numerous adaptations to withstand the constant barrage of both biotic and abiotic stressors. Occasionally, an adaptation emerges that may incidentally act as a virulence factor when an environmental microbe encounters a susceptible host (3–5). In contrast to host-acquired pathogens, environmentally acquired pathogens seemingly do not require a host stage in their life cycles and display virulence traits shaped by selection in a distinct ecological niche (5). For example, the mechanisms that protect *Legionella* and *Salmonella* strains from protozoan soil predators also aid their survival within human macrophages (6, 7). The production of Shiga toxins by *Escherichia coli* O157:H7 to fend off grazing protozoa (*Tetrahymena pyriformis*) is thought to aid colonization of humans (8). The opportunistic fungal pathogen *Rhizopus microsporus* is likewise endowed with resistance to both amoebae and human macrophages by its bacterial endosymbiont (*Ralstonia pickettii*), illustrating how an environmentally evolved tripartite interaction can lead to enhanced virulence(9).

A prime example of a bacterium equipped with a variety of competitive traits for survival in its ecological niche is *Chromobacterium haemolyticum* (10–13). First isolated from sputum (10), numerous strains have subsequently been isolated from freshwater sources (11–13). This facultatively anaerobic, Gram-negative bacterium is known for its pronounced antibacterial effect against *Salmonella*, *E. coli*, *Listeria monocytogenes*, and *Staphylococcus aureus* (14). In addition, *C. haemolyticum* displays mosquitocidal activity and inhibits the growth of *Plasmodium falciparum* through the production of cyanide and the histone deacetylase inhibitor romidepsin, respectively (13, 15). In recent years, there has been an alarming increase in the number of reports of severe opportunistic infections in animals (16) and humans (17–24) caused by *C. haemolyticum*. In many cases, *C. haemolyticum* infections are associated with exposure to freshwater, indicating a potential transmission route from the natural habitat to the host (16, 18–21, 24).

*C. haemolyticum* can infect different organs and body parts causing proctocolitis (17), pneumonia (18, 19), meningitis (20), or soft tissue infections (21, 22) that can lead to bacteremia (21, 23) and eventually sepsis (23, 24). These infections are difficult to treat, due to a combination of resistance of *C. haemolyticum* to antibiotics (10, 25, 26) and its frequent misidentification as *C. violaceum*, the more common representative of the *Chromobacterium* genus, potentially leading to ineffective treatment (18–20, 23). One of the most striking characteristics of *C. haemolyticum* is its pronounced β-haemolytic activity, which gives the species its name (10). Haemolysis can serve as a key virulence factor in mammalian infections, particularly given its association with sepsis (27). Although various virulence factors of *C. haemolyticum* have been linked to human (28, 29) and animal (13–15) disease, the haemolytic phenotype of *C. haemolyticum* has not yet been investigated in this regard, and its molecular basis remains unknown.

Here, we report the analysis of a human pathogenic *C. haemolyticum* strain by comparative genomics and metabolic profiling. We characterize a family of cyclic lipodepsipeptides produced by a non-ribosomal peptide synthetase (NRPS), which we find are responsible for *in vitro* haemolysis. Additionally, we demonstrate a critical role for the cyclic lipodepsipeptides in swarming and biofilm formation of *C. haemolyticum*. Taken together, our findings present a rare case of dual-use specialized metabolites that can both promote niche colonization and presumably contribute to the virulence of an environmentally acquired pathogen.

## Results

### Genome mining reveals a cryptic NRPS biosynthetic gene cluster that is conserved in human pathogenic *C. haemolyticum* strains

Because haemolysis can be induced by structurally diverse molecules, ranging from small peptidic natural products to large enzymes, we used a comparative genomics approach to try pinpoint the biosynthetic origin of haemolysins within the *Chromobacterium* genus. We noted a cryptic biosynthetic gene cluster (BGC) that is conserved among all *C. haemolyticum* strains, including the sputum isolate that prompted the recognition of the species, i.e. *C. haemolyticum* DSM 19808 (Fig. 1A). This putative BGC was named *chl* (76.8 kb, Fig. 1B). In this case of *C. haemolyticum* DSM 19808, the *chl* BGC was found on a contig edge of the publicly available whole genome sequence, necessitating the re- sequencing of the region (GenBank accession of *chl* BGC: 3793). Besides *C. haemolyticum*, only the phylogenetically closely related *Chromobacterium rhizoryzae* (30) and *Chromobacterium alkanivorans* (31) as well as four *Chromobacterium* sp. isolates harbor the *chl* BGC (Fig. 1A). This putative BGC has not yet been associated with a specific natural product but putatively codes for the biosynthesis of a nonribosomal peptide.

**Fig 1.**
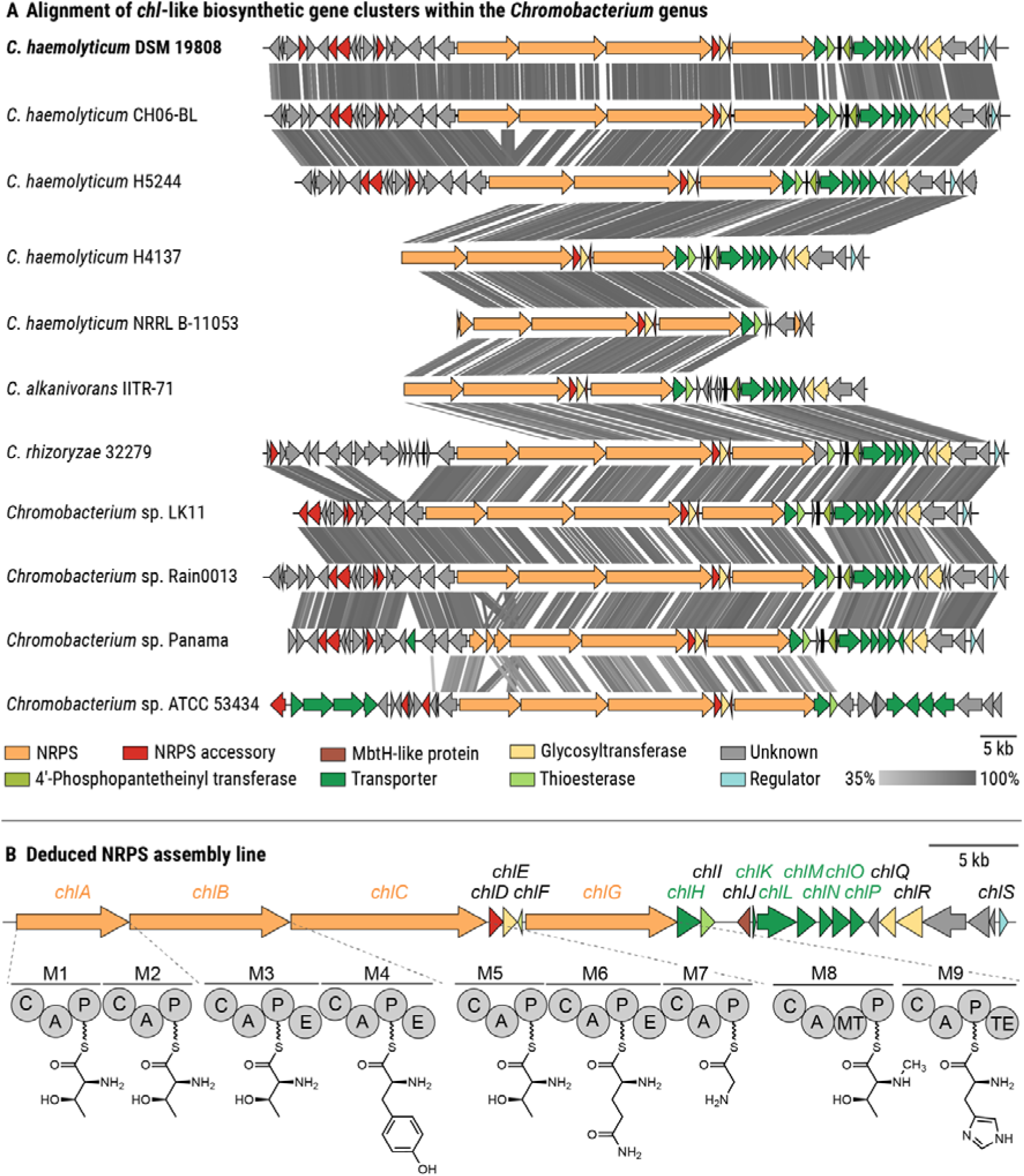
Identification of a cryptic NRPS-encoding biosynthetic gene cluster (BGC) in the genome of the opportunistic pathogen *C. haemolyticum*. (A) Alignment of *chl*-like gene clusters within the *Chromobacterium* genus. Colored arrows represent open reading frames, the putative products of which are indicated in the legend. The gray bar indicates the similarity of homologous regions, which are connected by gray lines. **(B)** Depiction of the deduced NRPS assembly line of the predicted *chl* BGC identified in the genome of *C. haemolyticum* (GenBank accession: 3793). Abbreviations: C: condensation domain; A: adenylation domain; P: peptidyl-carrier protein domain; E: epimerization domain, MT: *N*-methyltransferase domain; TE: thioesterase domain; M1 – M9: Module 1 – 9.

To determine whether the *chl* gene locus is unique to *Chromobacterium* spp., we constructed a genome neighborhood network using the ESI-Genome Neighborhood Tool and found similar BGCs in several cyanobacteria (e.g., *Anabaena* spp., *Cylindrospermopsis raciborskii*, and *Planktothrix sertra*) (Fig S1). These BGCs encode the biosynthetic enzymes for the production of hassallidins, which are cyclic lipopeptides with antifungal properties (32–35). We noted similarities in terms of gene assignments between the *chl* BGC and the *jag* BGC from the mushroom soft rot pathogen *Janthinobacterium agaricidamnosum* (Fig S1). Intriguingly, the corresponding secondary metabolite, jagaricin, is a cyclic lipopeptide with antifungal and haemolytic activities (36, 37).

Bioinformatic analysis of the *C. haemolyticum chl* BGC (38) revealed that the predicted NRPS is encoded by four genes (*chlA–C and chlG*), comprising nine modules responsible for chain elongation and an off-loading thioesterase (TE) domain (Fig 1B). The amino acid sequence of the nonribosomal peptide backbone was predicted using the Stachelhaus codes of the adenylation (A) domains, resulting in the tentative nonapeptide Thr–Thr–D-Thr–D-Tyr–Thr–D-Gln–Gly–*N*-Me-Thr–Leu (Fig 1B, Table S1). Noting that threonine residues can be dehydrated to the corresponding dehydrobutyrine (Dhb) residue via specialized condensation (C) domains (39), the peptide sequence resembles the core sequence found in both hassallidins and jagaricin, differing only in the predicted leucine at the C-terminal position (Table S1). Module 1 contains an additional condensation domain that could catalyze an *N-*terminal acylation, e.g., with a fatty acid. Due to the epimerization (E) domain present in module 3, 4, and 6, the corresponding Thr, Tyr, and Gln residues are predicted to be isomerized to the D-enantiomer. Module 8 contains an *N*-methyltransferase domain that might catalyze *N*-methylation of the threonine incorporated at the eighth position. Several additional biosynthetic enzymes are encoded in the BGC: a type-II-thioesterase (*chlI*) that could be responsible for restoring the NRPS activity when a module is misloaded (40, 41) and three glycosyl transferases (*chlE, chlQ*, and *chlR*) that might glycosylate the NRPS product, thus forming a glycopeptide. *N*- methylation at the eighth amino acid as well as the incorporation of sugar moieties are modifications that are also present in the hassallidins (35). Based on comparative genomics and bioinformatic prediction, our data indicate that *C. haemolyticum* DSM 19808 might produce a nonribosomal nonapeptide resembling jagaricin, which is known to display haemolytic activity. Therefore, we deemed the *chl* locus worthy of further investigation.

### Gene inactivation of *chl* leads to loss of haemolytic activity in *C. haemolyticum*

Due to the similarities between the *chl* and *jag* BGC and given that jagaricin is a known haemolysin (37), we aimed to target the *chl* BGC for inactivation and test whether the haemolytic phenotype of *C. haemolyticum* is affected (10). Seeing as there are no reports of genetic modification of *C. haemolyticum*, we adapted a double-crossover strategy that was developed for the related betaproteobacterium *J. agaricidamnosum* (36, 42). Using this strategy, the region encoding the C domain of module 4 in *chlB* was disrupted in the genome of *C. haemolyticum* DSM 19808 to generate *C. haemolyticum* Δ*chl* (*C. haemolyticum* Δ*chl*::kan^R^) (Fig 2A). *C. haemolyticum* DSM 19808 and *C. haemolyticum* Δ*chl* were inoculated on sheep blood agar and, after one day of incubation, their haemolytic activities were assessed. Wild-type *C. haemolyticum* forms a visible zone of haemolysis (Fig 2B), whereas *C. haemolyticum* Δ*chl* is devoid of haemolytic activity (Fig 2B). This observation indicates that the metabolites encoded by the *chl* BGC are behind the haemolytic activity of *C. haemolyticum*.

**Fig 2.**
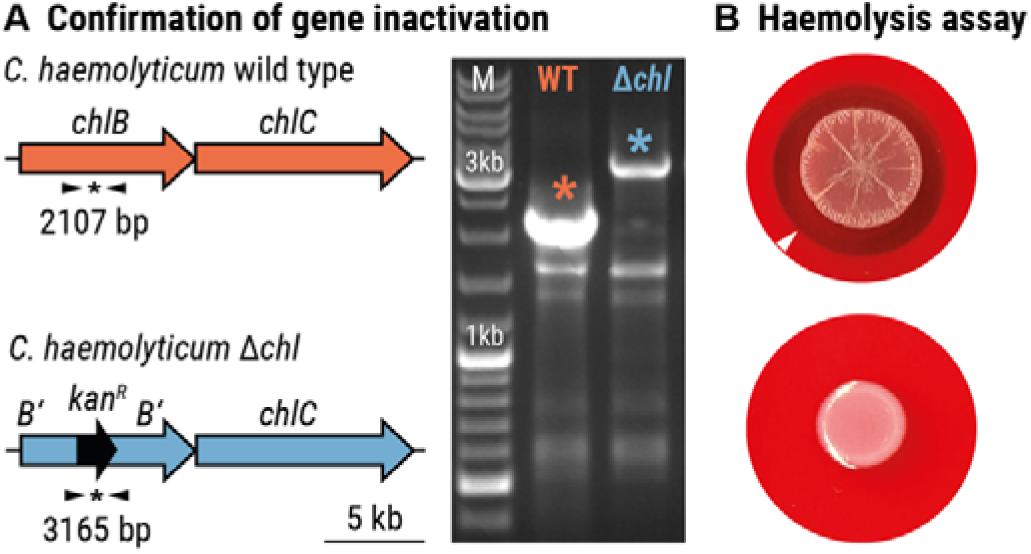
The products of the *chl*-encoded biosynthetic pathway mediate haemolytic activity of *C. haemolyticum*. **(A)** Verification of gene inactivation resulting from homologous recombination and incorporation of an antibiotic resistant cassette. Left: primer binding sites (black arrows) and expected size of amplicons (indicated by an *). Right: agarose gel with amplicons obtained by PCR of genomic DNA of *C. haemolyticum* wild type (wt) and *C. haemolyticum* Δ*chl*. **(B)** Colonies on sheep blood agar of *C. haemolyticum* wild type surrounded by a transparent zone of lysed erythrocytes (top panel) and *C. haemolyticum* Δ*chl*, which is not haemolytic (bottom panel). The white arrow head indicates the boundary of the zone of haemolysis.

### The *chl* gene locus codes for the production of the cyclic lipodepsipeptides jagaricin, chromolysin A and chromolysin B

With an intact *chl* BGC implicated in the haemolytic phenotype of *C. haemolyticum* DSM 19808, we sought to identify the cognate NRPS product by metabolic profiling. To this end, we cultivated *C. haemolyticum* DSM 19808 in a variety of media and analyzed the culture extracts via high-performance liquid chromatography coupled with high-resolution mass spectrometry (HPLC/HRMS). We detected a number of species sharing an MS² fragmentation pattern that indicates a peptide sequence in line with our bioinformatic prediction. A molecular network based on MS² data was constructed to fully explore the quantity and structural similarity of putative congeners, revealing 37 structurally related metabolites (Fig 3A). As none of these compounds are produced by *C. haemolyticum* Δ*chl*, we conclude that the *chl* gene locus codes for the biosynthesis of these 37 congeners (Fig S2).

**Fig 3.**
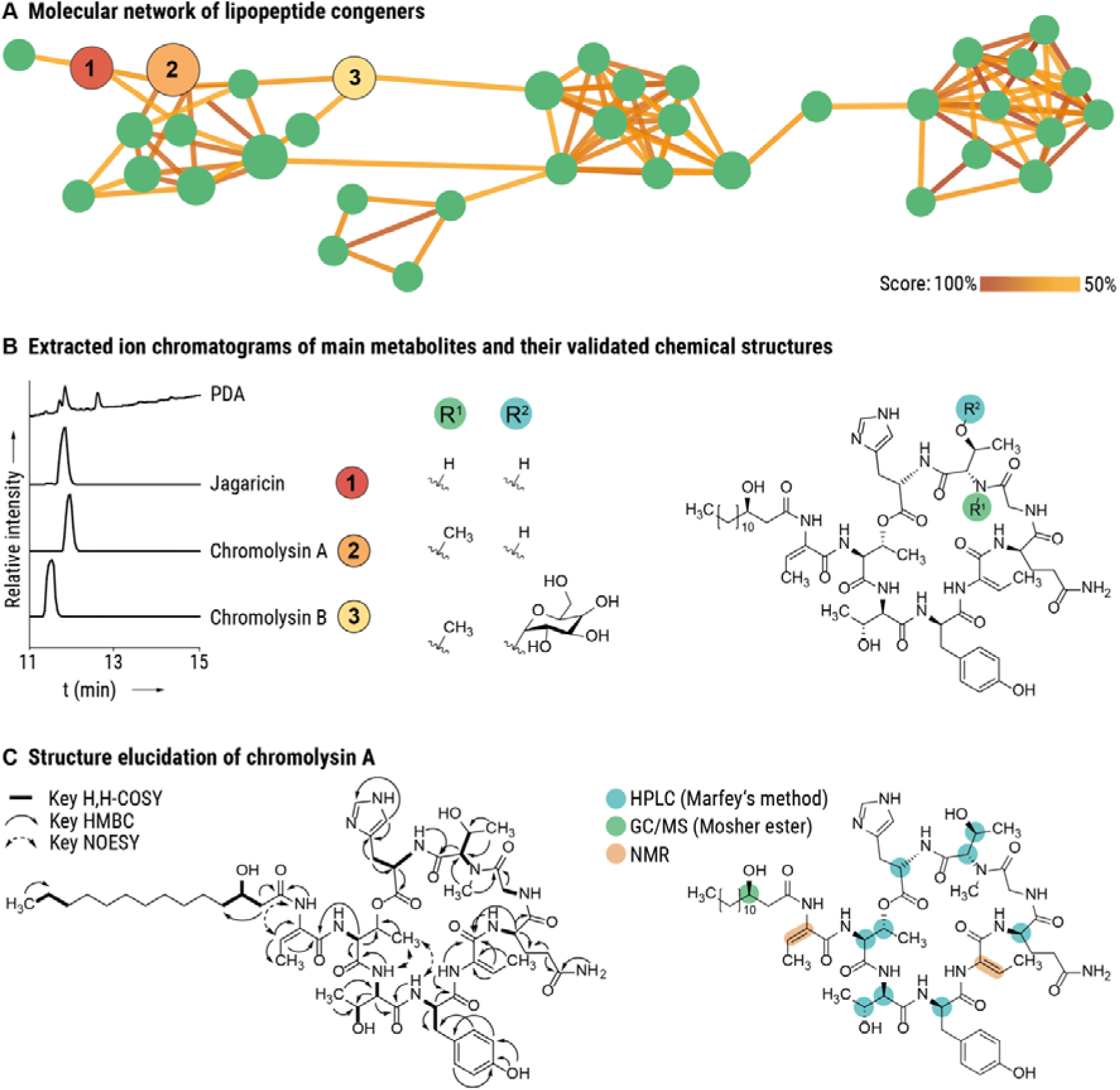
Discovery and elucidation of cyclic lipodepsipeptide metabolites. **(A)** Molecular network revealing entirety of lipopeptide congeners. Each node represents one compound; colored lines indicate structural similarity between two nodes based on MS² fragmentation. **(B)** Metabolic profile of *C. haemolyticum* DSM 19808 showing extracted ion chromatograms corresponding to the main metabolites and their validated chemical structures (jagaricin, chromolysin A and B). **(C)** Structure elucidation of chromolysin A. Left: Key 2D NMR couplings; Right: Assignment of absolute configuration.

The metabolic profiles showed that three main congeners were produced under the tested cultivation conditions (Fig 3B). The following sum formulas were deduced from HRMS: **1**: C_56_H_85_N_12_O_16_ (*m*/*z* 1181.6202 [*M*+H]^+^, calc. 1181.6201); **2**: C_57_H_87_N_12_O_16_ (*m*/*z* 1195.6359 [*M*+H]^+^, calc.

1195.6358); and **3**: C_63_H_97_N_12_O_21_ (*m*/*z* 1357.6906 [*M*+H]^+^, calc. 1357.6886). We noted that the sum formula of **1** matches that of jagaricin. Indeed, comparison of MS^2^ spectra and HPLC retention time with a jagaricin standard confirmed that *C. haemolyticum* produces jagaricin (**1**) (Fig S2). Congeners **2** and **3** were named chromolysin A and chromolysin B, respectively.

In order to obtain sufficient amounts of pure compounds for structural elucidation and bioactivity tests, a total of 15 L culture was extracted. Compound isolation using size-exclusion chromatography followed by preparative reverse-phase HPLC (RP-HPLC) yielded 0.6 mg of jagaricin, 5.6 mg of chromolysin A, and 1.5 mg of chromolysin B. We elucidated the structure of chromolysin A using nuclear magnetic resonance (NMR) spectroscopy and determined the absolute configuration by derivatization followed by HPLC or gas chromatography coupled with mass spectrometry (GC/MS) analysis. These analyses revealed that chromolysin A is an *N*-methylated congener of jagaricin (Fig 3C, Fig S3, Table S1).

The deduced sum formula of chromolysin B deviates from that of chromolysin A by C_6_H_10_O_5_ and shows an MS² fragmentation pattern that differs by a neutral loss of a C_6_H_10_O_5_ fragment (Fig S3). This indicates the presence of a sugar monomer in chromolysin B, which is likely incorporated by one of the glycosyl transferases encoded in the *chl* BGC. Based on MS²-fragmentation, we assume that *O*-glycosylation happens at the *N*-Me-Thr moiety, as is the case for the closely related hassallidin family (35) We isolated three additional congeners in sub-milligram amounts. These compounds putatively diverge by the length of the fatty acid chains and a replacement of histidine by tryptophan, as determined by NMR and MS²-fragmentation (Fig S4 – S8). Chromolysin A and B are identical to the previously described compounds Sch 20561 and Sch 20562, respectively, which are produced by human pathogenic *Aeromonas* species (43, 44) and putatively by mosquitocidal *Chromobacterium* sp. Panama, a strain that harbors a BGC similar to *chl* (Table S1) (15). A compound analogous to chromolysin A, named chromobactomycin, is also produced by the root-colonizing *Chromobacterium* sp. strain C61. Although chromolysin A and chromobactomycin have identical sum formulas, they differ in the order of the amino acids (45). In summary, we have identified the products of the *chl* BGC as a suite of cyclic lipodepsipeptides.

### Chromolysin A and B are potent haemolytic agents

In order to assess the potencies of the three main cyclic lipodepsipeptides produced by *C. haemolyticum* DSM 19808, we tested jagaricin, chromolysin A, and chromolysin B in a liquid erythrocyte lysis assay. Sheep erythrocytes were incubated with dilutions of each compound and haemolysis was quantified by measuring the released haemoglobin in the supernatants. All three compounds display potent haemolytic activity in the low micromolar range (jagaricin: 50 % haemolytic concentration [HC_50_] = 1.66 µM, 95 % confidence interval [CI] = 1.56–1.78 µM; chromolysin A: HC_50_ = 2.1 µM, CI = 1.96–2.23 µM; and chromolysin B: HC_50_ = 4.26 µM, CI = 4.19–4.33 µM; Fig 4A), which is comparable with the previously reported potency of jagaricin towards human erythrocytes (HC_50_ = 3.52 µM) (37). Our results demonstrate that, like jagaricin, chromolysin A and chromolysin B lyse erythrocytes.

**Fig 4.**
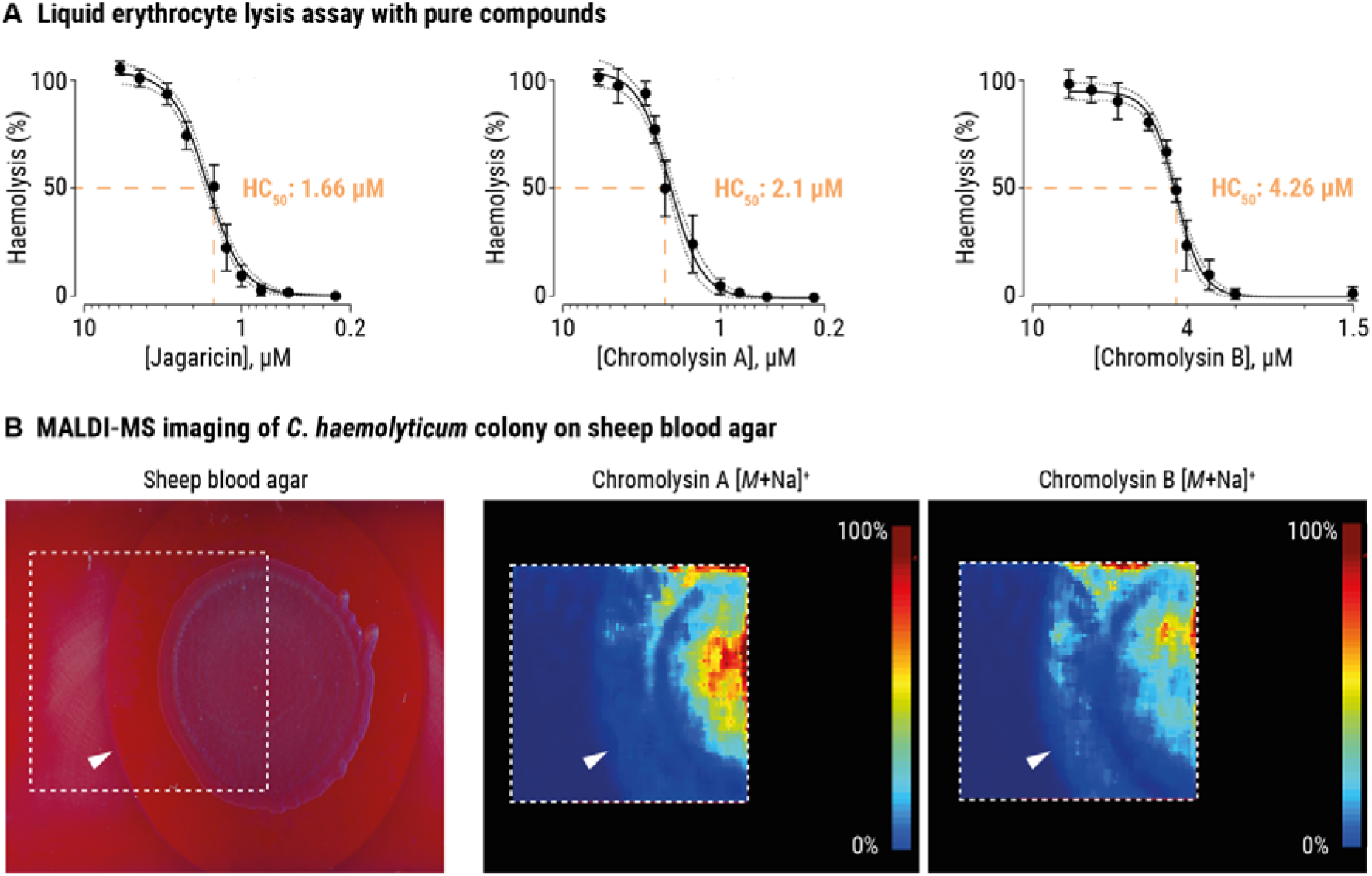
Chromolysins lyse sheep erythrocytes. **(A)** Haemolysis curve and HC_50_ values of jagaricin (left), chromolysin A (center), and chromolysin B (right). Data points represent the averages derived from four independent experiments (n = 4 biological replicates, with three technical replicates each) and the error bars represent the standard deviation (± S.D.). **(B)** MALDI-MS imaging of *C. haemolyticum* grown on sheep blood agar (left image). The distribution of chromolysin A (middle image) and chromolysin B (right image) in the colony and haemolytic zone is indicated by a MALDI-MS imaging heat map, in which the color-coded score (0–100 %) indicates the abundance of ion *m*/*z* = 1217.6 Da (+/– 0.5 Da, chromolysin A, [*M*+Na]^+^) and ion *m*/*z* = 1379.6 Da (+/– 0.5 Da, chromolysin B, [*M*+Na]^+^). Color code: blue = low abundance, red = high abundance of ion. The white arrow head indicates the boundary of the zone of haemolysis and the dashed square indicates the field of view for MALDI-MS imaging analysis.

To visualize the spatial distribution of the cyclic lipodepsipeptides within and around a *C. haemolyticum* colony, we used matrix-assisted laser desorption ionization mass spectrometry (MALDI-MS) imaging. We cultured *C. haemolyticum* DSM 19808 on sheep blood agar until a clear zone of haemolysis appeared surrounding the colony. The bacterial colony and zone were sprayed with a liquid matrix, and the imaging area was scanned. We detected only chromolysin A and B, not jagaricin, within the colony and in the haemolytic plaque (Fig 4B).

### Chromolysin A and B promote swarming and biofilm formation

Since cyclic lipopeptides often function as biosurfactants, which can promote biofilm formation and swarming motility (46–51), we investigated whether *chl*-derived lipodepsipeptides contribute to these important processes. To test whether the presence of an intact *chl* BGC is essential for biofilm formation, we cultivated the *C. haemolyticum* wild type and the *C. haemolyticum* Δ*chl* mutant in liquid cultures under slow agitation and visually monitored biofilm formation. After five days, wild-type *C. haemolyticum* produces a thick biofilm at the liquid-air interface, whereas *C. haemolyticum* Δ*chl* lacks the ability to form adhesive aggregates (Fig 5A).

**Fig 5.**
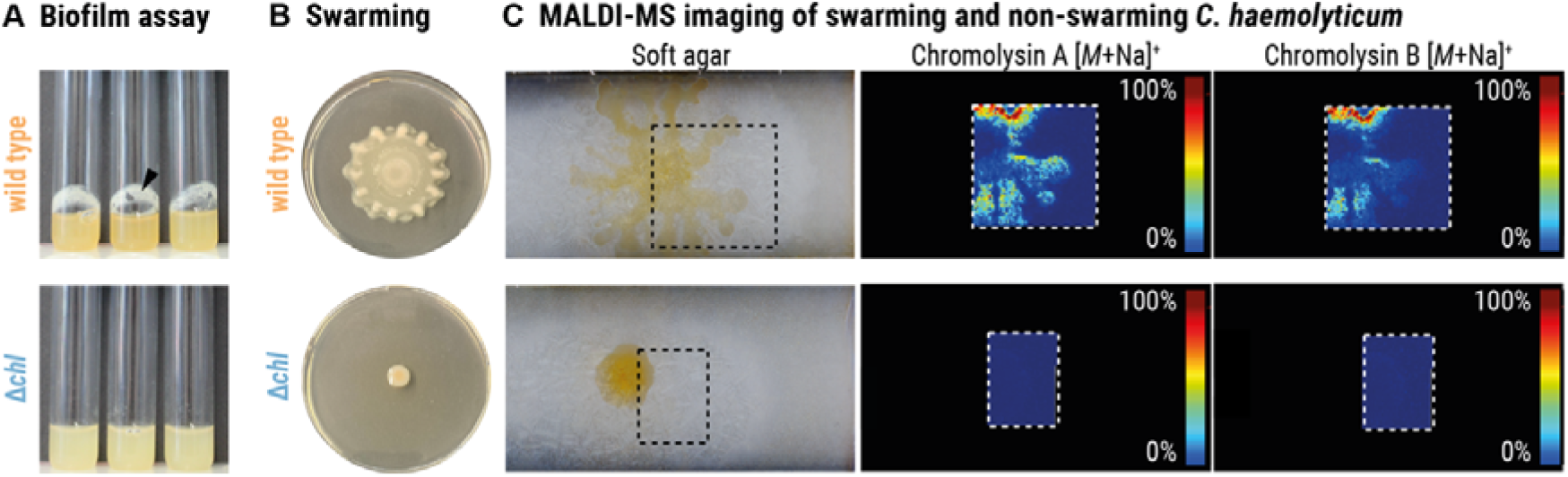
Chromolysin A and B promote swarming and biofilm formation. **(A)** Biofilm assays of *C. haemolyticum* wild type (top) and *C. haemolyticum* Δ*chl* (bottom) grown in liquid cultures. The black arrow head indicates the location of adhesive aggregates. **(B)** Swarming behavior of *C. haemolyticum* wild type (top) and *C. haemolyticum* Δ*chl* (bottom) on soft agar. **(C)** MALDI-MS imaging of *C. haemolyticum* wild type (top) and *C. haemolyticum* Δ*chl* (bottom) on soft agar demonstrates the accumulation of chromolysin A and chromolysin B in the dendritic formations of the wild type. The color-coded score (0–100 %) indicates the abundance of ion *m*/*z* = 1217.6 Da (+/– 0.5 Da, chromolysin A, [*M*+Na]^+^) and ion *m*/*z* = 1379.6 Da (+/– 0.5 Da, chromolysin B, [*M*+Na]^+^). Color code: blue = low abundance, red = high abundance of ion. The dashed square indicates the field of view for MALDI-MS imaging analysis.

We cultured *C. haemolyticum* wild type and *C. haemolyticum* Δ*chl* on soft agar for 24 hours to investigate their swarming ability. *C. haemolyticum* wild type radiates outwards in dendritic formations to colonize a large area of the agar surface, whereas *C. haemolyticum* Δ*chl* is unable to swarm (Fig 5B). We inspected the *C. haemolyticum* wild type colony with MALDI-MS imaging to visualize the distribution of jagaricin, chromolysin A, and chromolysin B during swarming. Of the three congeners tested, only chromolysin A and B accumulate in the dendritic formations of the swarming *C. haemolyticum* wild type colony, suggesting enhanced secretion in the periphery to improve motility. As expected, no ions corresponding to jagaricin, chromolysin A, and chromolysin B were detected in the case of *C. haemolyticum* Δ*chl* (Fig 5C). The impaired ability of chromolysin-deficient *C. haemolyticum* to swarm and to produce biofilms provides evidence that chromolysins are crucial for both processes, which may assist *C. haemolyticum* in colonizing new ecological niches (41, 49).

## Discussion

Most ecosystems encompass a remarkable diversity of microbes and higher organisms that compete with one another for space and resources. One bacterial species that is evidently primed for such inter-organismal competition is *C. haemolyticum*, which is antagonistic towards coinhabitants of its freshwater niche including bacteria (14), insects (13), and protists (15). *C. haemolyticum* is also equipped to transition to a pathogenic lifestyle, capable of infecting both humans (17–24) and other mammals (16). Despite mounting reports of serious disease caused by *C. haemolyticum*, the molecular mechanisms underlying pathogenicity are largely unknown (28). In this study, we reveal a family of cyclic lipodepsipeptides that are responsible for the characteristic β-haemolytic phenotype of *C. haemolyticum*. Furthermore, we implicate these specialized metabolites in the swarming and biofilm-forming behavior of *C. haemolyticum*, which may impact niche colonization.

By means of comparative genomics, we discovered a cryptic NRPS-encoding BGC (*chl*) that is universally conserved in *C. haemolyticum* strains. We noted the putative BGC to bear striking similarity to that of the jagaricin, a cyclic lipodepsipeptide with reported haemolytic activity (37). Accordingly, we found that *C. haemolyticum* Δ*chl*, in which the *chl* BGC has been disrupted, lacks haemolytic activity. Using a combination of metabolic profiling, tandem mass spectrometry, and NMR, we went on to demonstrate that *C. haemolyticum* also produces jagaricin, as well as a large number of closely related congeners, and that these specialized metabolites originate from the *chl* BGC. The haemolytic potencies of the three major congeners (jagaricin, chromolysin A and chromolysin B) (1.66–4.26 µM) were quantified and found to be comparable to that previously reported for jagaricin (3.52 µM) (37), despite differences in some assay parameters (e.g., sheep vs. human blood, measurement of absorbance at 405 nm vs. 541 nm, respectively). The haemolytic activity of jagaricin, chromolysin A, and chromolysin B is likely due to their amphiphilic character, each consisting of a cyclic peptide moiety covalently linked to a lipid chain, which can cause disruption of membrane integrity (52). Perhaps surprisingly, given its status as an emerging opportunistic pathogen, the molecular basis of the haemolytic phenotype of *C. haemolyticum* was until now enigmatic. It should be mentioned that other haemolysins remain to be discovered within the *Chromobacterium* genus since several other species are weakly haemolytic, namely *C. paludis* (53), *C. violaceum* (54), and *C. rhizoryzae* (30), although only the latter harbors a *chl*-like BGC (72 % sequence similarity). Indeed, the haemolytic activity of *C. violaceum* is lost when proteins are inactivated through heat treatment or enzymatic digestion, indicating the likely involvement of a pore-forming protein (54).

Notably, jagaricin was first reported as a major contributor to lesion formation in soft rot disease of the white button mushroom *Agaricus bisporus*, which is caused by *J. agaricidamnosum* (36). Jagaricin was also found to display broad anti-eukaryotic effects, inhibiting numerous pathogenic fungi and social amoebae, as well as being toxic towards human cells including erythrocytes (36, 37) . Given that *J. agaricidamnosum* is not known to infect mammals, the biological relevance (if any) of the latter was unclear, and appeared to be incidental to its membrane-disrupting properties (37). Yet, it is likely that the haemolytic properties of jagaricin come to the fore during *C. haemolyticum* infection. Our results therefore exemplify an intriguing case of context-dependent bioactivity, in which the same specialized metabolite is utilized by taxonomically distant bacteria (*J. agaricidamnosum* and *C. haemolyticum* belong to different orders within the class Betaproteobacteria) to manifest disease in distinct eukaryotic hosts. It is also possible that the anti-fungal and amoebicidal effects of jagaricin (36, 37) provide both *J. agaricidamnosum* and *C. haemolyticum* with a competitive advantage in their respective environmental niches.

Regardless of structure, haemolysins are often associated with the pathogenic potential of their producers (55–57). One pertinent example is streptolysin S (SLS), a ribosomally synthesized and post-translationally modified peptide haemolysin that is an indispensable virulence factor of human pathogenic *Streptococcus pyogenes* (58). As is the case for many haemolysins, SLS destructs erythrocytes as a means to access nutrients, primarily iron (58–60). Iron is an essential element since it plays catalytic, regulatory, and structural roles, as well as being instrumental as a virulence regulator (61). Perhaps jagaricin, chromolysin A, and chromolysin B similarly participate in iron acquisition during the infection process of *C. haemolyticum*, thereby aiding survival in the host. Fittingly, the *chl* BGC harbours six genes (*chlK–P*) that encode homologs of proteins involved in the transport of haemin (62, 63). Based on the resemblance of ChlK–P to the periplasmic binding-protein-dependent transport system for haemin (HemPRSTUV) of the animal pathogen *Yersinia enterocolitica* (64, 65), one could envision the following process occurring in *C. haemolyticum*: ChlK and ChlL proteins transport haemin across the outer membrane into the periplasm, where it is bound by ChlN. ChlM degrades haemin in the periplasm to liberate iron, which is transported into the cytoplasm across the inner membrane by ChlO and ChlP (64, 65).

Given that lipopeptides often act as biosurfactants that contribute to the swarming and biofilm-forming behavior of their producers, thereby facilitating movement along surfaces (66) and access to new niches (46–51), we probed a similar role for the products of the *chl* BGC of *C. haemolyticum*. Indeed, *C. haemolyticum* Δ*chl* is deficient in swarming and biofilm formation. Our observations are in accordance with previous reports of cyclic lipopeptides positively influencing swarming and biofilm formation (50, 67). It is conceivable that both processes contribute to the survival of *C. haemolyticum* not only in the environment, but also during the host stage. Swarming properties might equip *C. haemolyticum* with the ability to colonize animals if it encounters a susceptible host, then biofilm formation might enable subversion of innate immune defenses during the infection process (68). As previously mentioned, the haemolytic activity of the cyclic lipodepsipeptides could cause erythrocyte lysis, leading to iron release and accompanying enhanced bacterial proliferation. Notably, haemolysins can also be cytolytic for cells of the immune system, e.g., macrophages and neutrophils, which could further support immune evasion (58, 69). We thus propose that the *chl*-derived cyclic lipodepsipeptides could be considered “dual-use” virulence factors, a concept that refers to the observation that factors that promote persistence in the environment can also promote virulence within a host (70), ultimately resulting in “accidental virulence” (71). This fits with the well-documented opportunistic nature of *C. haemolyticum* infection. Lastly, it is worth mentioning the possible translational avenues arising from the data presented herein. The enzymatic cleavage of cyclic lipopeptides is commonly employed by competing bacteria in the environment (72, 73) and is considered a promising alternative approach to treat bacterial infections (74, 75). Thus, the products of the *chl* BGC may represent promising targets for the development of antivirulence therapeutics to combat *C. haemolyticum* infections.

Despite its increasing recognition as an opportunistic pathogen, little is currently known regarding the features that shape the pathobiology of *C. haemolyticum*. Similarly, there is limited understanding of genetic determinants of niche colonization by *C. haemolyticum*. Here, we provide insight into both aspects by deeming jagaricin, chromolysin A, and chromolysin B as lynchpin chemical mediators not only responsible for the pronounced haemolytic phenotype of *C. haemolyticum* but crucial for biofilm formation and swarming motility. Our experimental evidence points to the possibility that *C. haemolyticum* employs dual-use specialized metabolites to survive and thrive in its environmental niche, while simultaneously being able to successfully switch to a pathogenic lifestyle once inside a human host. In illuminating the virulence mechanisms of *C. haemolyticum*, we lay the foundation for the development of antivirulence strategies against this serious emerging pathogen.

## Materials and Methods Strains and growth conditions

*Chromobacterium haemolyticum* DSM 19808 (MDA0585^T^) was maintained on Lysogeny Broth (LB) (10 g/L tryptone, 5 g/L yeast extract, 5 g/L NaCl, 1 g/L glucose, 15 g/L agar) at 37 °C. For metabolic screening, bacteria were cultured in Medium 2 (2 g/L yeast extract, 0.45 g/L KH_2_PO_4_, 2.39 g/L Na_2_HPO_4_·12H_2_O, 1 g/L beef extract, 5 g/L NaCl, 5 g/L peptone), Rich Medium (5 g/L yeast extract, 10 g/L peptone, 5 g/L casamino acids, 2 g/L beef extract, 5 g/L malt extract, 2 g/L glycerin, 1 g/L MgSO_4_·7H_2_O, 0.05 g/L Tween 80), Jagaricin Production Medium (20 g/L D-mannitol, 20 g/L D-fructose, 10 g/L yeast extract, 5 g/L sodium L-glutamate, 6 mL/L trace element solution (4 g/L CaCl_2_·H_2_O, 1 g/L C_6_H_5_FeO_7_·H_2_O, 0.2 g/L MnSO_4_, 0.1 g/L ZnCl_2_, 0.04 g/L CuSO_4_·5H_2_O, 0.03 g/L CoCl_2_·6H_2_O, 0.03 g/L Na_2_MoO_4_·2H_2_O, 0.1 g/L Na_2_B_4_O_7_·10H_2_O)) (76), MGY (1.25 g/L yeast extract, 10 g/L glycerol, 7 g/L K_2_HPO_4_, 2 g/L KH_2_PO_4_, 0.588 g/L sodium citrate, 1 g/L (NH_4_)_2_SO_4_, 0.1 g/L MgSO_4_), LB (10 g/L tryptone, 5 g/L yeast extract, 5 g/L NaCl, 1 g/L glucose), or potato dextrose broth (4 g/L potato extract, 20 g/L glucose). *Escherichia coli* TOP10 (Invitrogen, Thermo Fisher Scientific) and DNA-methyltransferase deficient *E. coli* ER2925 (New England Biolabs) were used for construction and amplification of plasmids and cultured on solid LB medium at 37 °C. *C. haemolyticum* mutants were selected on solid LB medium supplemented with 50 µg/mL kanamycin.

## Bioinformatic analysis

BGCs resembling the *chl* BGC were identified in bacterial genomes of the NCBI database using the EFI Genome Neighborhood Tool (EFI-GNT) (77). Using the integrated BLASTp tool, products of the NRPS-encoding genes (GJA_RS00800–GJA_RS00815) were compared to protein sequences of the UniProt database (BLAST sequences ≤ 100; e-value ≤ 10^−5^; neighborhood window size 10). Results were analyzed using antiSMASH 6.0 (78). A synteny diagram (e-value ≤ 10^−3^; length ≥ 100; identity value ≥ 35) was created using EasyFig 2.2.4 (79) and the integrated tBLASTx tool, and was recolored with Adobe Illustrator.

## Generation of metabolic profiles

Bacterial cultures (100 mL) were grown for three to four days, then extracted with 100 mL ethyl acetate. The organic layers were dried over anhydrous Na_2_SO_4_, filtered using Macherey-Nagel MN 615 ¼ filter paper and concentrated under reduced pressure. The crude extracts were dissolved in 1.3 mL methanol, of which 25 µL was diluted with 25 µL methanol before analysis by HPLC/HRMS using an Exactive Hybrid-Quadrupole-Orbitrap with electrospray ion source coupled to an Accela HPLC system (Thermo Fisher Scientific). For separation, a Betasil C18 column (2.1 × 150 mm, 3 μm, Thermo Fisher Scientific) was used with gradient elution (solvent A: H_2_O + 0.1 % HCOOH, solvent B: MeCN + 0.1 % HCOOH, gradient: 0–1 min 5 % B, 1–16 min 5–98 % B, 16–20 min 98 % B4; flow rate 0.2 mL/min).

## Tandem mass spectrometry

Samples were dissolved in methanol, of which 3 µL was injected and analyzed using a QExactive Hybrid-Quadrupole-Orbitrap with electrospray ion source coupled to an Accela HPLC system (Thermo Fisher Scientific). For separation, an Accucore C18 column (2.1 × 100 mm, 2.6 μm, Thermo Fisher Scientific) was used with gradient elution (solvent A: H_2_O + 0.1 % HCOOH, solvent B: MeCN + 0.1 % HCOOH, gradient: 0–10 min 5–98 % B, 10–14 min 98 % B; flow rate 0.2 mL/min). MS^2^ spectra were generated for molecular network analysis by the top 5 analysis method (scan range *m*/*z* 400–1500, 25 % normalized collision energy (HCD)).

## Mass spectrometry network analysis

A molecular network was created with MS^2^ spectra from *C. haemolyticum* culture extracts using Compound Discoverer 3.3. Only metabolites that yielded fragment ions were chosen for cluster generation. Two nodes were linked when the following parameters were fulfilled: Msn Score ≥ 50 %, minimal coverage ≥ 60 %, matching fragments ≥ 10. Node size is proportional to the peak area of the respective compound in the LC/MS spectrum. The colors of the links depend on the Msn Score (50– 100 %) of the two linked metabolites. The network was subsequently cropped and recolored using Adobe Illustrator.

## Production and isolation of lipopeptides

*C. haemolyticum* was grown in 30 × 500 mL volumes of LB medium, totaling 15 L, for three days at 30 °C without agitation. Each 500 mL culture was extracted with 300 mL of ethyl acetate and the combined organic layers were dried over anhydrous Na_2_SO_4_, filtered and concentrated under reduced pressure. The crude extract was fractionated by size exclusion chromatography (Sephadex LH-20 column (440 × 35 mm) with methanol as eluent). Fractions containing the desired compounds were further purified by preparative HPLC (Shimadzu LC-8a series, DAD detector) using a C-18 Grom- Saphir 110 column (250 × 20 mm, 5 µm) and gradient elution (solvent A: H_2_O + 0.01 % TFA, solvent B: MeCN, gradient: 3 min 10 % B, 10–90 % B in 35 min; flow rate 10 mL/min). Fractions containing each of jagaricin, chromolysin A, and chromolysin B were collected, then separately combined and lyophilized to yield 0.6 mg, 5.6, and 1.5 mg purified compound, respectively. For physicochemical data, see Supporting Information.

## Nuclear magnetic resonance (NMR) spectroscopy

NMR spectra were recorded in deuterated dimethyl sulfoxide (DMSO-*d_6_*) using a Bruker AVANCE III 600 MHz instrument equipped with Bruker Cryo Platform. Chemical shifts are reported in ppm relative to the solvent residual signal (^1^H: δ = 2.50 ppm, ^13^C: δ = 39.52 ppm) (80). The following abbreviations are used for multiplicities of resonance signals: s = singlet, d = doublet, t = triplet, q = quartet, br = broad.

## Synthesis of *N*-methyl threonine reference compounds

Synthesis was performed as previously described (81, 82) with the following modifications. SOCl_2_ (910 µL) was added dropwise to dry methanol (3.5 mL), and the solution was cooled to –10 °C. L-Thr (350 mg) was then added slowly, and the reaction mixture was stirred for 14 h at ambient temperature before being diluted with 2 mL of a saturated aqueous NaHCO_3_ solution. The mixture was extracted with CH_2_Cl_2_ (four times); the combined organic extracts were dried over anhydrous Na_2_SO_4_, filtered using Macherey-Nagel MN 615 ¼ filter paper and concentrated under reduced pressure yielding the threonine methyl ester (L-Thr-O-Me). L-Thr-O-Me (110 mg) was dissolved in CH_2_Cl_2_ (11 mL) and 0.1 N aqueous trifluoroacetic acid (TFA, 10 mL) was added before cooling the mixture to 0 °C. Formaldehyde (37 % aqueous solution, 69 µL) was added dropwise under vigorous stirring, which was continued for 8 h at ambient temperature before being neutralized with a saturated aqueous NaHCO_3_ solution. The mixture was extracted with CH_2_Cl_2_ (four times). The combined organic extracts were dried over anhydrous Na_2_SO_4_, filtered using Macherey-Nagel MN 615 ¼ filter paper, and concentrated under reduced pressure yielding the corresponding oxazolidine derivative, which was used without further purification. Oxazolidine (8 mg) was dissolved in dry CH_2_Cl_2_ (0.8 mL), cooled to 0 °C, and TFA (0.8 mL) was added followed by dropwise addition of triethyl silane (0.09 mL). The reaction mixture was stirred at ambient temperature for 24 h before being concentrated under reduced pressure. The pale-yellow residue was resuspended in 1 N HCl and washed with petroleum ether. Hydrolysis in 6 N HCl (1 mL) under reflux for 16 h yielded 8 mg *N*-Me-L-Thr hydrochloride salt as a pale-yellow foam. The same procedure was used to synthesize *N*-Me-L-*allo*-Thr, *N*-Me-D-Thr and *N*-Me-D-*allo*-Thr from L-*allo*-Thr, L-Thr and L-*allo*-Thr as staring material, respectively.

## Absolute configuration of amino acids

Hydrolysis of 1 mg chromolysin A was carried out in 1 mL of 6 M aqueous HCl solution supplemented with 0.05 % phenol under reflux for 14 h. A 500 µL aliquot was concentrated under reduced pressure and then dissolved in 100 µL water and 50 µL 1 M aqueous NaHCO_3_ solution. Subsequently, 10 µL of a freshly prepared 10 mg/mL solution of 1-fluoro-2,4-dinitrophenyl-5-L-alanine-amide (L-FDAA) in acetone was added. The reaction mixture was stirred at 40 °C for 1 h before being quenched by addition of 25 µL 2 M aqueous HCl solution followed by 25 µL MeOH. Reference amino acids (1 mg) were treated identically and all samples were diluted 1:4 with MeOH before analysis via analytical HPLC (Shimadzu LC-10 Avp series, DAD detector, LiChrospher 100 RP-18 endcapped column (250 × 4.6 mm, 5 µm), flow rate 1 mL/min, solvent A: H_2_O + 0.1 % TFA, solvent B: MeCN).

For determination of the retention times of the synthetic references, two different gradients were applied: Gradient A (0–4 min 25 % B, 4–44 min 25–40 % B, 44–45 min 40–98 % B), provided the retention times [min] of D-His 6.19, L-His 7.94, L-*allo*-Thr 13.19, D-*allo*-Thr 14.59, *N*-Me-D-Thr 15.25, D-Thr 17.18, *N*-Me-L-*allo*-Thr 18.65, *N*-Me-D-*allo*-Thr 19.94, L-Tyr 25.39 and D-Tyr 28.88.

Gradient B (0–5 min 20 % B, 5–35 min 20–25 % B, 35–40 min 25–98 % B) yielded sufficient separation of D-Gln 22.81, L-Gln 23.34, L-Thr 23.87, *N*-Me-L-Thr 25.73 and D-*allo-*Thr 28.62.

The absolute configurations of the amino acid moieties were assigned as L-His (7.97 min, gradient A), L-Thr (23.87 min, gradient B), *N*-Me-L-*allo*-Thr (18.81 min, gradient A), D-*allo-*Thr (14.68 min, gradient A and 28.58 min, gradient B), D-Gln (22.80 min, gradient B) and D-Tyr (29.19 min, gradient A).

Configuration of dehydrobutyrine moieties was deduced from ^1^H-NMR chemical shifts of the vinylic proton quartets (δ *=* 5.82 and 5.63), which correlated with the reported ^1^H-NMR data of chromolysin A (Sch 20561, δ *=* 5.81 and 5.70) and its glycosylated congener chromolysin B (Sch 20562, δ *=* 5.84 and 5.80) (43, 44). By synthesis of the *N*-acetylated and *O*-methylated (*E*)- and (*Z*)- isomers, it was shown that the vinylic proton of the (*Z*)-isomer (δ *=* 6.48) is downshifted in comparison to the (*E*)-isomer (δ *=* 5.90) (44).

## Absolute configuration of **β**-hydroxy fatty acid

Derivatization of the free fatty acid was performed as described previously (83) with the following modifications. Hydrolyzed chromolysin A (0.5 mg) was dissolved in 200 µL MeOH and 30 µL of a 2 M solution of (trimethylsilyl)diazomethane in hexane were added to the reaction mixture. After stirring at ambient temperature for 10 min, the mixture was dried under nitrogen flow. The residue was dissolved in 400 µL of a 0.2 M solution of 4-dimethylaminopyridine in dichloromethane and 15 µL of (*R*)-(–)-α- methoxy-α-(trifluoromethyl)phenylacetyl chloride were added. The reaction mixture was stirred at ambient temperature for 2 h before being dried under nitrogen flow. During the reaction, a color change of the solution from colorless to golden-yellow was observed. Reference fatty acids β-(*R*)- hydroxymyristic acid (HMA) and β-(*R,S*)-HMA (1 mg) were treated identically and all samples were dissolved in MeOH before analysis via GC/MS.

GC/MS analysis was performed on a Trace 1310 GC (Thermo Scientific) coupled with a TSQ 9000 electron impact (EI)-triple quad mass spectrometer (Thermo Scientific). A 4 mm SSL GC inlet glass liner with glass wool (P/N 453A1305) and a BPX5 capillary column (30 m, 0.25 mm inner diameter, 0.25 μm film) from Trajan (SGE) were used. The column was operated using helium carrier gas (0.6 mL/min) and split injection (split flow: 25 mL/min, split ratio: 15). The temperature of the injector was set to 250 °C. The oven temperature was set to 40 °C for 4 min and was then increased to 120 °C with a rate of 40 °C/min followed by a temperature increase to 250 °C with a rate of 0.75 °C/min. The MS transfer line was set to 300 °C, the ion source temperature was set to 200 °C. Total ion current (TIC) values were recorded in the mass range of 45–500 amu with an offset of 70 min and a dwell time of 0.2 s. The injection volume was 0.5 μL.

The references and the probe show peaks with an identical fragmentation pattern containing the main fragment ions *m*/*z* 189.0 and *m*/*z* 241.2 with a retention time of 142.94 min (β-(*R*)-HMA); 142.51 min and 142.92 min (β-(*R,S*)-HMA) and 142.94 min (probe). The fatty acid was therefore assigned as β-(*R*)-HMA.

## Generation of *C. haemolyticum* Δ*chl* using homologous recombination

The target for genetic inactivation was the second open reading frame of the putative NRPS BGC (ctg20_66). Two neighboring homologous regions (hr1 and hr2, ∼940 bp) were amplified byPCR using the primer pair *nrps*_C4_hr1_fw/*nrps*_C4_hr1_rv / *nrps*_C4_hr2_fw/*nrps*_ C4_hr2_rv (Table S2). Both DNA regions were extended by 15 bp to be complementary to the *Sma*I-linearized vector pGL42a (84), and by 8 bp overlapping a kanamycin resistance cassette (*kan*^R^). *Kan*^R^ was amplified using the primer pair *nrps*_*kan*^R^_fw / *nrps*_*kan*^R^_rv (Table S2) with an 8 bp overhang that binds hr1 and hr2, respectively. Following agarose gel electrophoresis, the resulting amplicons (hr1, hr2, and *kan^R^*) were purified using the Monarch DNA Gel Extraction Kit (New England Biolabs) and subsequently assembled using the In-Fusion HD Cloning Kit and the *Sma*I-linearized cloning vector pGL42a. The resulting plasmid (pKO*nrps*) was introduced into chemically competent TOP10 *E. coli* (Invitrogen, Thermo Fisher Scientific) following the manufacturer’s instructions. Transformants were selected on LB agar supplemented with 50 µg/mL kanamycin. Incubation proceeded at ambient temperature until colonies formed, which were cultivated overnight in 2 mL LB medium at 37 °C with agitation. The vector pKO*nrps* was isolated using the Monarch Plasmid Miniprep Kit (New England Biolabs), then verified using restriction digest followed by Sanger sequencing. Subsequently, pKO*nrps* was transferred to DNA methylase-deficient *E. coli* ER2925 before being isolated to obtain unmethylated vector for transfer to *C. haemolyticum*. Competent *C. haemolyticum* cells were generated following a protocol previously developed for *Mycetohabitans rhizoxinica* (85); however, the growth medium was changed to LB. Unmethylated pKO*nrps* (20–100 ng) was added to 60 µL of cells prior to electroporation at 25 µF, 200 Ω and 2.5-3 kV (Eppendorf Eporator®, Eppendorf, Hamburg, Germany). The cells were recovered in 0.5 mL LB at 37 °C for 3–4 h with orbital shaking, and spread on LB agar supplemented with 50 µg/mL kanamycin. The resulting colonies were checked for integration of *kan^R^*at the targeted region of the genome by colony PCR using Δ*nrps* verification primers (Table S2). Once verified, *C. haemolyticum* Δ*chl* was conserved at –80 °C in 50 % glycerol cryogenic stocks.

## MALDI mass spectrometry imaging

To localize haemolytic natural products via imaging mass spectrometry, 10 µL of a *C. haemolyticum* or *C. haemolyticum* Δ*chl* overnight culture was pipetted onto tryptic soy agar supplemented with 5 % sheep blood (Oxoid) and cultivated for 24 h at ambient temperature.

To localize natural products involved in swarming, 10 µL of a *C. haemolyticum* or *C. haemolyticum* Δ*chl* overnight culture was pipetted onto MGY soft agar (0.6 % agar) and cultivated for 24 h at ambient temperature. Agar with bacterial growth was subsequently transferred to an indium tin oxide-coated slide and dried overnight at 37 °C. Dried samples were sprayed with a 1:1 mixture of 20 mg/mL 2,5-dihyroxybenzoic acid and α-cyano-4-hydroxycinnamic acid in MeCN:MeOH:H_2_O (70:25:5) on an ImagePrep Station 2.0 (Bruker Dynamics) in 60 consecutive cycles (spraying for 1 s, incubating for 10 s, drying for 10 s). After 30 cycles, samples were rotated by 180° to ensure even coverage.

The samples were analyzed withan UltrafleXtreme MALDI TOF/TOF (Bruker Daltonics) in positive reflector mode with flexControl 3.0. The flexControl method was calibrated with the Peptide Calibration Standard II (Bruker) before every measurement. Each sample was irradiated with 500 laser impulses in a 200 µm wide grid (10 random impulses per grid position), and ions in a mass range of *m*/*z* 160–2160 were detected. The resulting spectra were processed with FlexImaging 4.1 or SCiLs™ Lab 2016b with baseline subtraction and analyzed for masses of interest (± 0.5 Da) that were subsequently visualized. For graphical documentation, imaging heat maps were cropped and the contrast was adjusted using Adobe Illustrator.

## Biofilm formation

LB medium (2 mL) was inoculated with *C. haemolyticum* DSM 19808 or *C. haemolyticum* Δ*chl* and incubated at 30 °C with slow agitation (120 rpm). After five days, cultures were observed for the occurrence of biofilm on the glass wall.

## Erythrocyte lysis assay

A 1 mL volume of defibrinated sheep erythrocytes (Thermo Scientific Oxoid) was washed by resuspension in 50 mL phosphate buffered saline (PBS) (137 mM NaCl, 2.7 mM KCl, 10 mM Na_2_ HPO_4_, 2 mM KH_2_PO_4_; pH 7.4), followed by centrifugation at 300 × *g* for 10 min at 20 °C. The supernatant, containing haemoglobin released from lysed erythrocytes during storage at 4 °C, was gently decanted and the erythrocytes were washed a second time. The intact erythrocytes remaining after washing were resuspended in PBS to give an approximately 3–4 % *v/v* suspension of erythrocytes in PBS. The erythrocyte suspension was adjusted with PBS so that the absorbance reading obtained when treated with the nonionic surfactant Triton X-100, which is commonly used as a positive control in haemolysis assays, was within the linear range at 405 nm (see below).

The erythrocyte lysis assay (86) was performed by adding 75 μL volumes of the erythrocyte suspension to wells of a 96-well U-bottom microtiter plate (PS, clear; Greiner Bio-One). 75 μL volumes of all control and test solutions were added in triplicate (technical replicates) to erythrocyte-containing wells (1:1 dilution), resulting in a final erythrocyte concentration of approximately 1.5–2 % *v/v*. The positive control (corresponding to 100 % or complete haemolysis) solution was 0.2 % Triton X-100 (0.1 % *v/v* Triton X-100 final concentration). The vehicle control as (corresponding to 0 % or background haemolysis) solution was PBS containing 4 % DMSO (2 % *v/v* DMSO final concentration). Purified **2**, **3** and **4** were dissolved in PBS containing 4 % DMSO (2 % *v/v* DMSO final concentration) to give test solutions covering a range of compound concentrations.

After incubation at 37 °C for 1 h, the microtiter plate was centrifuged at 300 × *g* for 5 min at 20 °C and 100 μL volumes of the supernatants were transferred to a fresh microtiter plate using a pipette, ensuring not to disturb any sedimented intact erythrocytes. Haemoglobin release was measured by measuring the absorption of the supernatants at 405 nm (A405) using a Varioskan LUX 3020-237 (Thermo Scientific) microplate reader with SkanIt Software for Microplate Readers (Research Edition, Product Version 7.0.0.50; Thermo Fisher Scientific).

The absorbance values from each set of technical replicates were averaged and the extent of haemolysis (percentage haemolysis) caused by each compound concentration was calculated using the formula: [(average A405 of the treated samples at a given concentration – average A405 of vehicle-treated samples) / (average A405 of Triton-treated samples – average A405 of vehicle-treated samples)] × 100. The entire experiment was performed four times on different days (biological replicates). The normalized percentage haemolysis values were plotted against the logarithmic value (log_10_) of the corresponding compound concentrations (µM) tested to determine the HC_50_ values using GraphPad Prism version 10.1.2 for Windows (GraphPad Software, Boston, Massachusetts USA; www.graphpad.com). The data points represent the average and the error bars represent the standard deviation (±S.D.) derived from four independent experiments (biological replicates; n=4). The best fit dose-response curve was determined using a log(inhibitor) vs. response; variable slope (four parameters) fit. The fitting method was least squares regression, with no weighting and with the bottom parameter fixed to zero. Confidence intervals (95 %, profile-likelihood) are shown.

## Data availability

All data generated or analyzed during this study are included in the manuscript and in the supporting files.

## Acknowledgments

Financial support by the Deutsche Forschungsgemeinschaft (DFG, German Research Foundation) under Germany’s Excellence Strategy – EXC 2051 (Cluster of Excellence ’Balance of the Microverse’) Project-ID 390713860 and the SFB 1127 ChemBioSys, Project-ID 239748522, and the Leibniz Award (to C.H.) is gratefully acknowledged. We thank Sarah P. Niehs for helpful discussions regarding the design of analytic experiments.

## Author Contributions

L.D., P.W., I.R., K.S., and C. H. conceived the idea and developed the study design. L.D., P.W., I.R., and E.M.M. interpreted the data. L.D. and P.W. performed bioinformatic analysis, bioassays, cultivation, metabolic profiling, and isolation of compounds. L.D., P.W., and K.S. elucidated structures.

P.W. generated mutant strains. E.M.M. performed the haemolysis microtiter assay. F.T. performed MALDI-imaging measurements. P.M. performed GC/MS measurements and method optimization.

L.M.J. performed DNA sequencing. S.J.P. performed DNA sequence assembly and analysis. L.D., I.R., E.M.M., and C. H. wrote the manuscript.

