## Supplementary File for "Dual-use virulence factors of the opportunistic pathogen *Chromobacterium haemolyticum* mediate haemolysis and colonization"

Supplementary file 1 for

**Dual-use virulence factors of the opportunistic pathogen *Chromobacterium haemolyticum* mediate haemolysis and colonization**

Leo Dumjahn^a,^*, Philipp Wein^a,^*, Evelyn M. Molloy^a^, Kirstin Scherlach^a^, Felix Trottmann^a^, Philippe Meisinger^a^, Louise M. Judd^b^, Sacha J. Pidot^b^, Timothy P. Stinear^b^, Ingrid Richter^a,#^, Christian Hertweck^a,c,d, #^

* These authors contributed equally to this work.

^#^ Co-corresponding authors: Ingrid Richter:; Christian Hertweck:

**Table S1.** Bioinformatic prediction (antiSMASH) of A domain specificity of a putative NRPS encoded by the *chl* biosynthetic gene cluster (BGC) of *Chromobacterium haemolyticum* DSM 19808 and *chl*-like BGCs. NCBI accession numbers are given for whole genome assemblies.

| **Name** | **A1** | **A2** | **A3** | **A4** | **A5** | **A6** | **A7** | **A8** | **A9** | **Genome assembly ID** |
| --- | --- | --- | --- | --- | --- | --- | --- | --- | --- | --- |
| *Chromobacterium haemolyticum* DSM 19808 | Thr | Thr | Thr | Tyr | Thr | Gln | Gly | Thr | Leu | [ASM71188v1](https://www.ncbi.nlm.nih.gov/datasets/genome/GCF_000711885.1/) |
| *Chromobacterium haemolyticum* CH06-BL | Thr | Thr | Thr | Tyr | Thr | Gln | Gly | Thr | Leu | [ASM993615v1](https://www.ncbi.nlm.nih.gov/datasets/genome/GCF_009936155.1/) |
| *Chromobacterium haemolyticum* UGAL515B_03 | Thr | Thr | Thr | Tyr | Thr | Gln | Gly | Thr | Leu | [ASM3309584v1](https://www.ncbi.nlm.nih.gov/datasets/genome/GCF_033095845.1/) |
| *Chromobacterium haemolyticum* Bb2 | Thr | Thr | Thr | Tyr | Thr | Gln | Gly | Thr | Leu | [ASM1478898v1](https://www.ncbi.nlm.nih.gov/datasets/genome/GCF_014788985.1/) |
| *Chromobacterium haemolyticum* H5244 | - | - | Thr | - | Thr | Gln | Gly | Thr | Leu | [ASM208185v1](https://www.ncbi.nlm.nih.gov/datasets/genome/GCF_002081855.1/) |
| *Chromobacterium haemolyticum* JD-14 | - | - | - | Tyr | Thr | Gln | Gly | Thr | Leu | [ASM1731578v1](https://www.ncbi.nlm.nih.gov/datasets/genome/GCF_017315785.1/) |
| *Chromobacterium haemolyticum* H4137 | - | - | - | Tyr | Thr | Gln | Gly | Thr | Leu | [ASM208182v1](https://www.ncbi.nlm.nih.gov/datasets/genome/GCF_002081825.1/) |
| *Chromobacterium haemolyticum* WJ-5 | - | - | - | Tyr | Thr | Gln | Gly | Thr | Leu | [ASM1731580v1](https://www.ncbi.nlm.nih.gov/datasets/genome/GCF_017315805.1/) |
| *Chromobacterium haemolyticum* ZR-1 | - | - | - | Tyr | Thr | Gln | Gly | Thr | Leu | [ASM1652242v1](https://www.ncbi.nlm.nih.gov/datasets/genome/GCF_016522425.1/) |
| *Chromobacterium haemolyticum* GD04138 | - | - | - | Tyr | Thr | Gln | Gly | Thr | Leu | [ASM2983525v1](https://www.ncbi.nlm.nih.gov/datasets/genome/GCF_029835255.1/) |
| *Chromobacterium haemolyticum* H3973 | - | - | - | Tyr | Thr | Gln | Gly | Thr | Leu | [ASM208181v1](https://www.ncbi.nlm.nih.gov/datasets/genome/GCF_002081815.1/) |
| *Chromobacterium haemolyticum* IR17 | - | - | - | Tyr | Thr | Gln | Gly | Thr | Leu | [ASM333214v1](https://www.ncbi.nlm.nih.gov/datasets/genome/GCF_003332145.1/) |
| *Chromobacterium haemolyticum* T124 | - | - | - | Tyr | Thr | Gln | Gly | Thr | Leu | [ASM75847v1](https://www.ncbi.nlm.nih.gov/datasets/genome/GCF_000758475.1/) |
| *Chromobacterium haemolyticum* NRRL B-11053 | - | - | - | Tyr | Thr | Gln | Gly | Thr | Leu | [ASM305254v1](https://www.ncbi.nlm.nih.gov/datasets/genome/GCF_003052545.1/) |
| *Chromobacterium rhizoryzae* 32279 | Thr | Thr | Thr | Tyr | Thr | Gln | Gly | Thr | Leu/His | [ASM2054446v1](https://www.ncbi.nlm.nih.gov/datasets/genome/GCF_020544465.1/) |
| *Chromobacterium alkanivorans* IITR-71 | - | - | - | Tyr | Thr | Gln | Gly | Thr | Leu | [ASM1693765v1](https://www.ncbi.nlm.nih.gov/datasets/genome/GCF_016937655.1/) |
| *Chromobacterium* sp. LK11 | - | - | Thr | Tyr | Thr | Gln | Gly | Thr | Leu | [ASM104370v1](https://www.ncbi.nlm.nih.gov/datasets/genome/GCF_001043705.1/) |
| *Chromobacterium* sp. Rain0013 | Thr | Thr | Thr | Tyr | Thr | Gln | Gly | Thr | Leu | [ASM1529176v1](https://www.ncbi.nlm.nih.gov/datasets/genome/GCF_015291765.1/) |
| *Chromobacterium* sp. Panama | - | - | Thr | Tyr | Thr | Gln | Gly | Thr | Leu | [ASM305255v1](https://www.ncbi.nlm.nih.gov/datasets/genome/GCF_003052555.1/) |
| *Chromobacterium* sp. ATCC 53434 | Thr | Thr | Thr | Tyr | Thr | Gln | Gly | Thr | Leu | [ASM284834v1](https://www.ncbi.nlm.nih.gov/datasets/genome/GCF_002848345.1/) |
| *Anabaena* sp. 90 | Thr | Thr | Thr | Tyr | Thr | Gln | Gly | Thr | Gln/Tyr | [ASM31270v1](https://www.ncbi.nlm.nih.gov/datasets/genome/GCF_000312705.1/) |
| *Cylindrospermopsis raciborskii* CS505 | Thr | Thr | Thr | Tyr | Thr | Gln | Gly | Thr | Tyr | [ASM167658v1](https://www.ncbi.nlm.nih.gov/datasets/genome/GCF_001676585.1/) |
| *Cryptosporiopsis* *curvispora* GIHE-G1 | Thr | Thr | Thr | Tyr | Thr | Gln | Gly | Thr | Gln/Tyr | [ASM1448941v1](https://www.ncbi.nlm.nih.gov/datasets/genome/GCF_014489415.1/) |
| *Raphidiopsis* *curvata* NIES-932 | Thr | Thr | Thr | Tyr | Thr | Gln | Gly | Thr | Gln/Tyr | [ASM236813v1](https://www.ncbi.nlm.nih.gov/datasets/genome/GCF_002368135.1/) |
| *Planktothrix serta* PCC 8927 | Thr | Thr | Thr | Tyr | Thr | Gln | Gly | Thr | Gln/Tyr | [PCC_8927_PRJEB10992_v2](https://www.ncbi.nlm.nih.gov/datasets/genome/GCF_900010725.2/) |
| *Calothrix brevissima* NIES-22 | Thr | Thr | Thr | Tyr | Thr | Gln | Gly | Thr | Gln/Tyr | [ASM236799v1](https://www.ncbi.nlm.nih.gov/datasets/genome/GCF_002367995.1/) |
| *Janthinobacterium* *agaricidamnosum* DSM 9628 | Thr | Thr | Thr | Tyr | Thr | Gln | Gly | Thr | His | [JAG1](https://www.ncbi.nlm.nih.gov/datasets/genome/GCF_000723165.1/) |

**Table S2.** Primers used for generating *C. haemolyticum* Δ*chl*. Non-binding overhangs are underlined.

| **Name** | **Oligo sequence 5’→3’** | **Size of amplicon (bp)** | **Purpose** |
| --- | --- | --- | --- |
| *nrps*_C4_hr1_fw | ACTAGTGGATCCCCCGAACAACTCAGCAGCGTGGTCAAC | 964 | pKO*nrps* construction |
| *nrps*_C4_hr1_rv | ATTCATCCGAGTCGAACAACTGGGCGTTGTC |  |  |
| *nrps*_*kan*^R^_fw | CCAGCAGCTCAGAAGAACTCGTCAAGAAGGCGATAG | 1074 |  |
| *nrps*_*kan*^R^_rv | TTCGACTCGGATGAATGTCAGCTACTGGGCTATCTGG |  |  |
| *nrps*_C4_hr2_fw | TCTTCTGAGCTGCTGGTCTTCGAGAACTACC | 967 |  |
| *nrps*_C4_hr2_rv | GAATTCCTGCAGCCCCCATGAAGATTTCCGACACCGAGG |  |  |
| *nrps*_KOver_out_fw | CGTCTTGATCCAACGCCG | 2588 | Δ*chl* verification |
| *nrps*_KOver_in_rv | GTCGTTACTTTTCCGGGCTG |  |  |
| *nrps*_KOver_in_fw | ACCTGGAACACTTCGCATTG | 2950 |  |
| *nrps*_KOver_out_rv | GATCTCTATCCGCTGTCCCC |  |  |

**Table S3.** NMR shifts of chromolysin A (**2**).

| **Partial structure** | **Position** | **^13^C NMR** | **^1^H NMR (mult., *J* in Hz)** |
| --- | --- | --- | --- |
| β-(*R*)-Hydroxymyristic acid |  |  |  |
|  | 1 | 170.5 | - |
|  | 2 | 43.7 | 2.24 (2H, m) |
|  | 3 | 67.4 | 3.78 (1H, m) |
|  | 4 | 37.0 | 1.33 (2H, m*) |
|  | 5-12 | 28.9-29.2 | 1.20-1.25 (16H, brs*) |
|  | 13 | 22.1 | 1.29-1.25 (2H, m*) |
|  | 14 | 14.0 | 0.85 (3H, t, 6.9) |
|  | 3-OH | - | 5.11 (1H, brd, 5.3) |
| (*E*)*-*Dehydrobutyrine |  |  |  |
|  | 1 | 164.4 | - |
|  | 2 | 130.2 | - |
|  | 3 | 121.5^+^ | 5.82 (1H, m) |
|  | 4 | 13.5 | 1.82 (3H, d, 7.3) |
|  | NH | - | 9.29 (1H, s) |
| l-Threonine |  |  |  |
|  | 1 | 168.7 | - |
|  | 2 | 55.5 | 4.62 (1H, m*) |
|  | 3 | 71.8 | 5.27 (1H, brd, 4.9) |
|  | 4 | 16.2 | 1.20 (3H, m*) |
|  | NH | - | 7.73 (1H, brs) |
| d-*allo*-Threonine |  |  |  |
|  | 1 | 169.4 | - |
|  | 2 | 58.8 | 4.17 (1H, m) |
|  | 3 | 66.7 | 3.92 (1H, m) |
|  | 4 | 20.4 | 1.07 (3H, d, 5.9) |
|  | 3-OH | - | 5.17 (1H, brs) |
|  | NH | - | 8.01 (1H, brd, 5.9) |
| d-Tyrosine |  |  |  |
|  | 1 | 170.2 | - |
|  | 2 | 54.7 | 4.48 (1H, dd, 13.6, 6.0) |
|  | 3 | 36.2 | 2.91 (1H, m*) |
|  |  |  | 2.77 (1H, m*) |
|  | 4 | 127.4 | - |
|  | 5, 9 | 130.0 | 6.94 (2H, d, 8.3) |
|  | 6, 8 | 114.9 | 6.59 (2H, d, 8.3) |
|  | 7 | 155.8 | - |
|  | 7-OH | - | 9.16 (1H, s) |
|  | NH | - | 8.32 (1H, m*) |
| (*E*)*-*Dehydrobutyrine |  |  |  |
|  | 1 | 164.7 | - |
|  | 2 | 131.5 | - |
|  | 3 | 118.8^+^ | 5.63 (1H, q, 7.1) |
|  | 4 | 13.0 | 1.78 (3H, d, 7.3) |
|  | NH | - | 9.51 (1H, s) |
| d-Glutamine |  |  |  |
|  | 1 | 171.2 | - |
|  | 2 | 52.5 | 4.32 (1H, m) |
|  | 3 | 27.1 | 2.07 (1H, m*) |
|  |  |  | 1.73 (1H, m*) |
|  | 4 | 31.5 | 2.14 (2H, m*) |
|  | 5 | 173.9 | - |
|  | 5-NH_2_ | - | 7.26 (1H, brs) |
|  |  |  | 6.78 (1H, brs) |
|  | NH | - | 8.21 (1H, m) |
| Glycine |  |  |  |
|  | 1 | 169.3 | - |
|  | 2 | 41.1 | 3.99 (2H, m) |
|  | NH | - | 7.96 (1H, brs) |
| *N*-Me-l-*allo*-Threonine | 1 | 169.1 | - |
|  | 2 | 62.1 | 4.63 (1H, m*) |
|  | 3 | 64.3 | 4.06 (1H, m) |
|  | 4 | 20.2 | 0.98 (3H, d, 5.9) |
|  | 3-OH | - | n. d. |
|  | N-CH_3_ | 31.3 | 2.81 (3H, s) |
| l-Histidine |  |  |  |
|  | 1 | 170.2 | - |
|  | 2 | 51.1 | 4.59 (1H, m*) |
|  | 3 | 25.0 | 3.08 (1H, m*) |
|  |  |  | 2.90 (1H, m*) |
|  | 4 | 129.8 | - |
|  | 5 | 134.0 | 8.78 (1H, brs) |
|  | 6 | 117.3 | 7.30 (1H, s) |
|  | 5-NH | - | 13.96 (1H, brs) |
|  | NH | - | 8.32 (1H, m*) |

Abbreviations: n. d.: not detected; *: partial overlap; ^+^: deduced using HSQC.

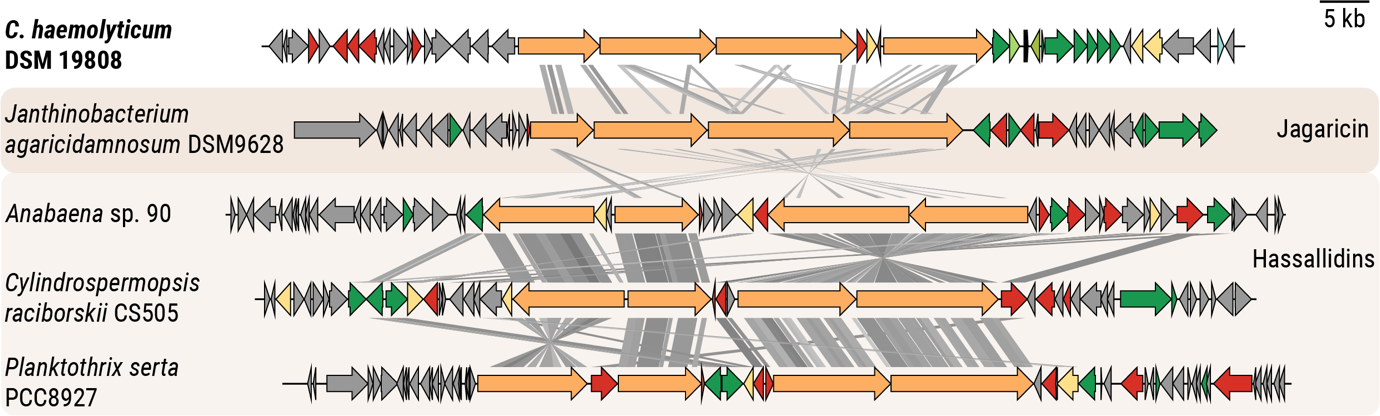

**Figure S1.** Alignment of the *chl* biosynthetic gene cluster (BGC) with similar BGCs from cyanobacteria and the soft rot pathogen *J.* *agaricidamnosum*.

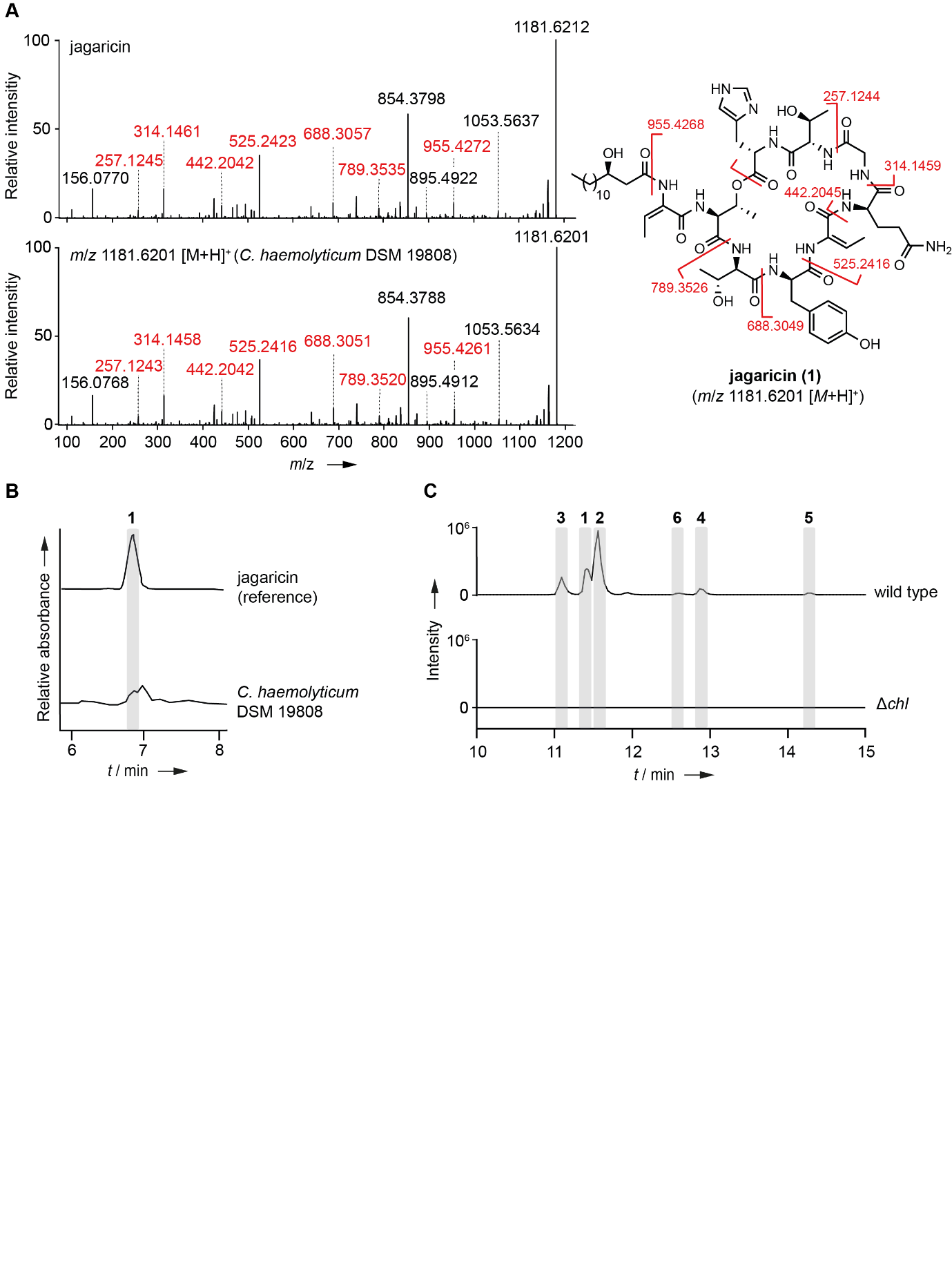

**Figure S2. Verification of the *chl* biosynthetic gene cluster of *C. haemolyticum* DSM 19808 as the genetic origin of jagaricin production.** **(A)** MS² fragmentation patterns of reference compound jagaricin (**1**) (top) and **1** detected in the metabolite profile of *C. haemolyticum* (bottom). Assignable fragments are colored in red; structure of **1** is depicted (right) with expected masses of assignable fragments given (calculated in ChemDraw 23.0.1). **(B)** HPLC profile of reference compound jagaricin (**1**) (top) compared to that of *C. haemolyticum* culture extract (bottom). **(C)** Extracted ion chromatogram of *m*/*z* values corresponding to [*M*+H]^+^ ions of jagaricin (**1**) and chromolysins A–E (**2**–**6**) in culture extracts of *C. haemolyticum* wild type and *C. haemolyticum* Δ*chl*.

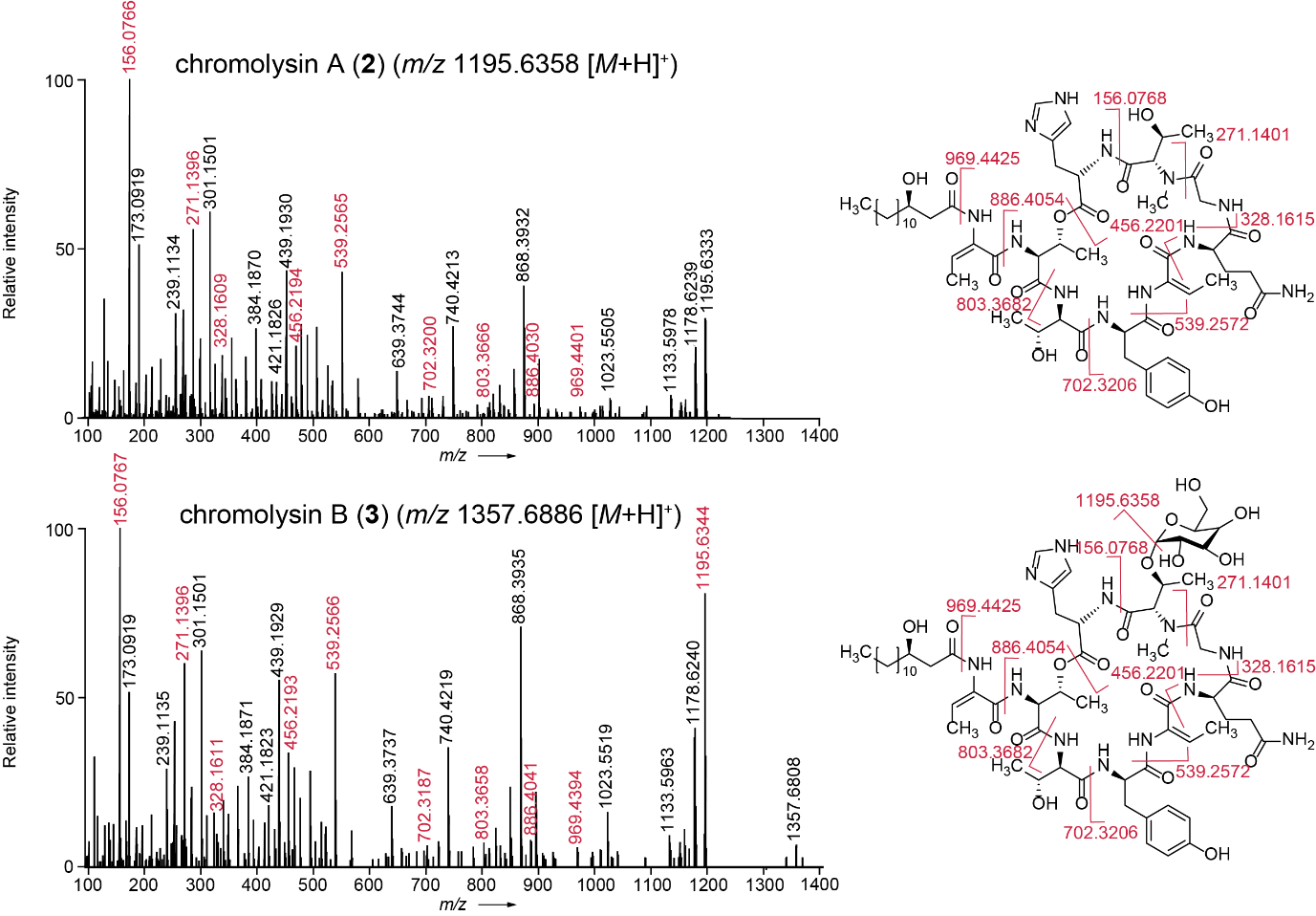

**Figure S3.** MS² fragmentation patterns of chromolysin A (**2**) and chromolysin B (**3**) in *C. haemolyticum* culture extracts. Assignable fragments are colored in red; respective structures (right) are depicted with expected masses of assignable fragments given (calculated in ChemDraw 23.0.1).

**A**

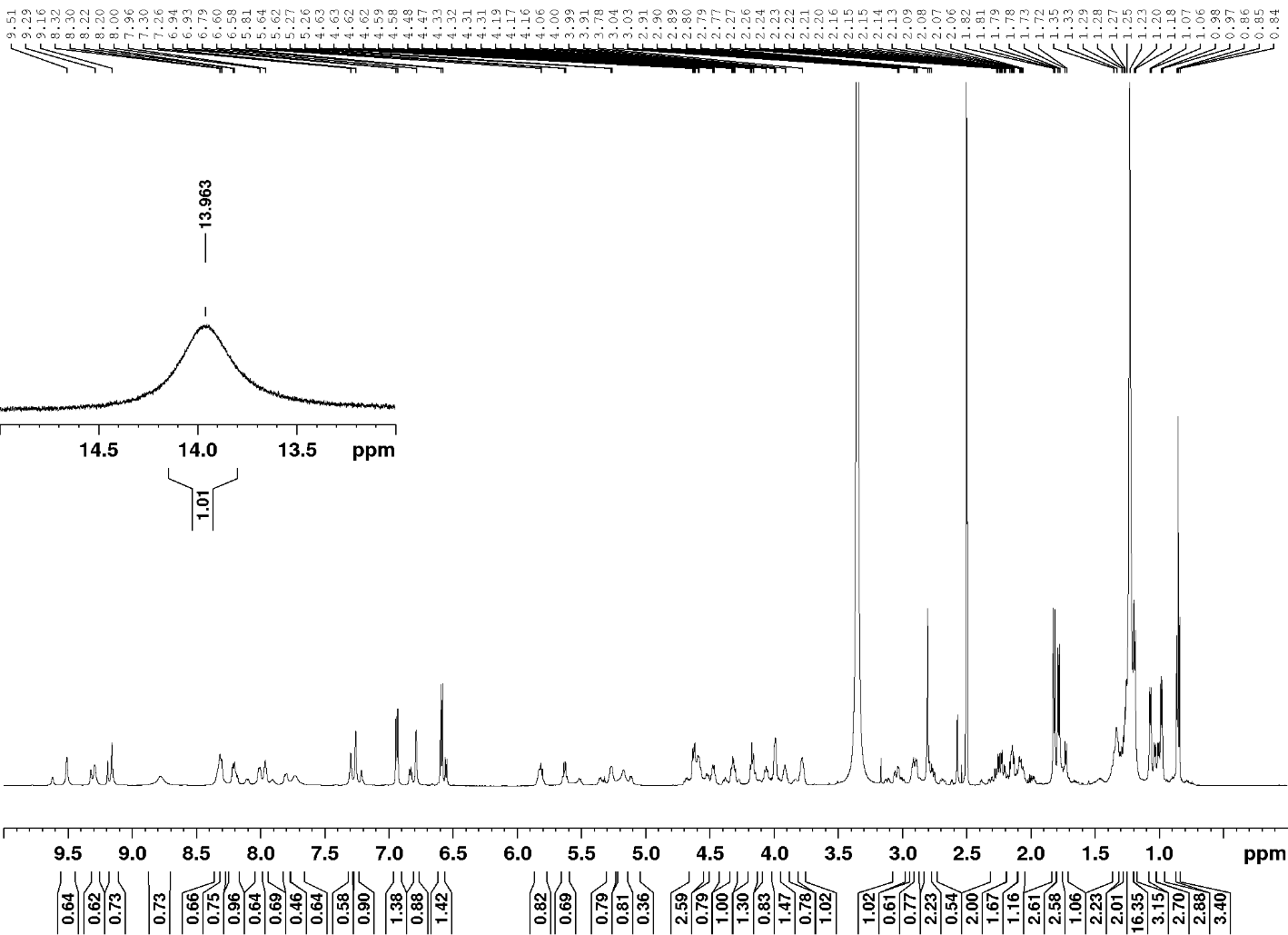

**B**

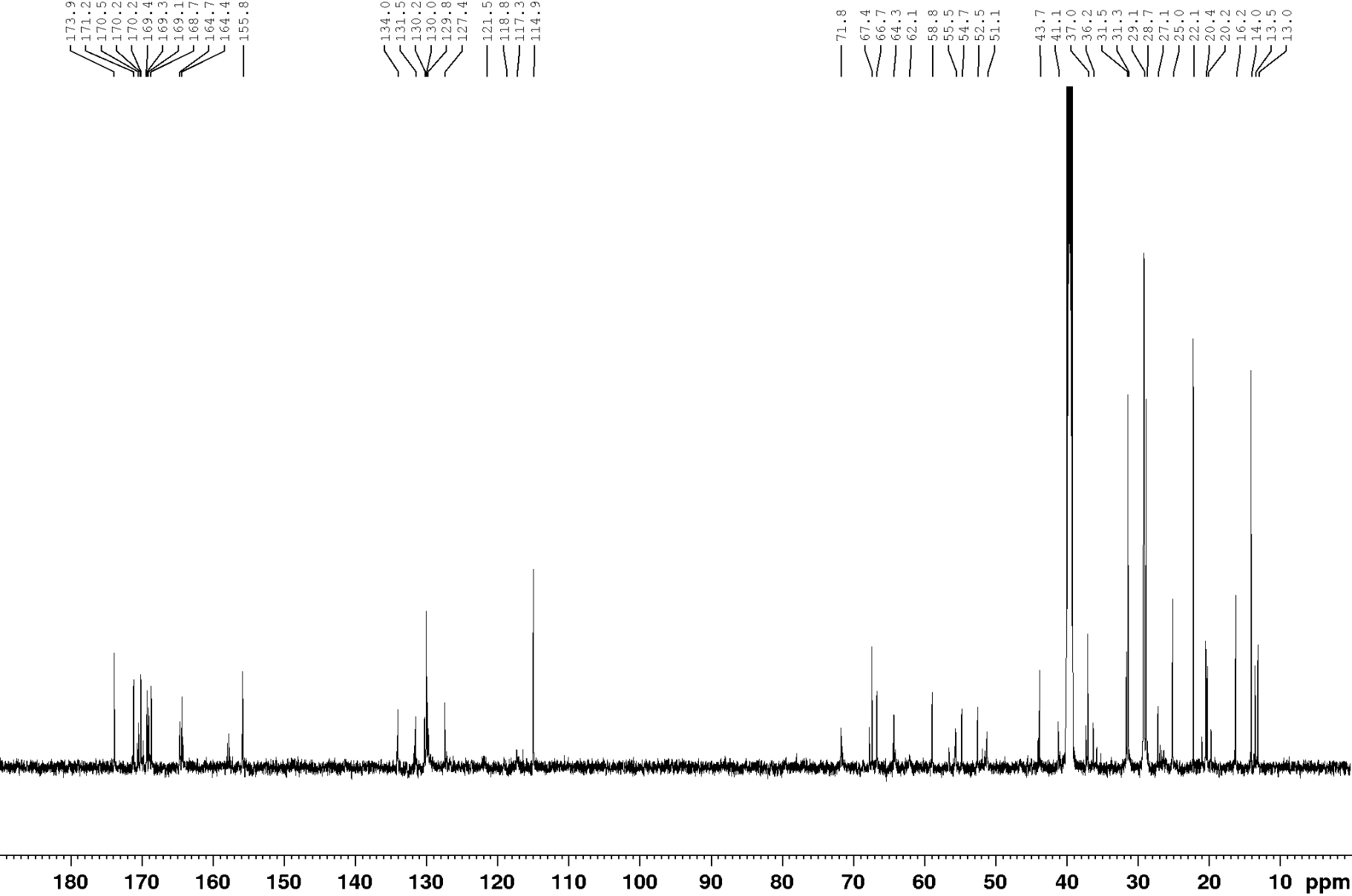

**C**

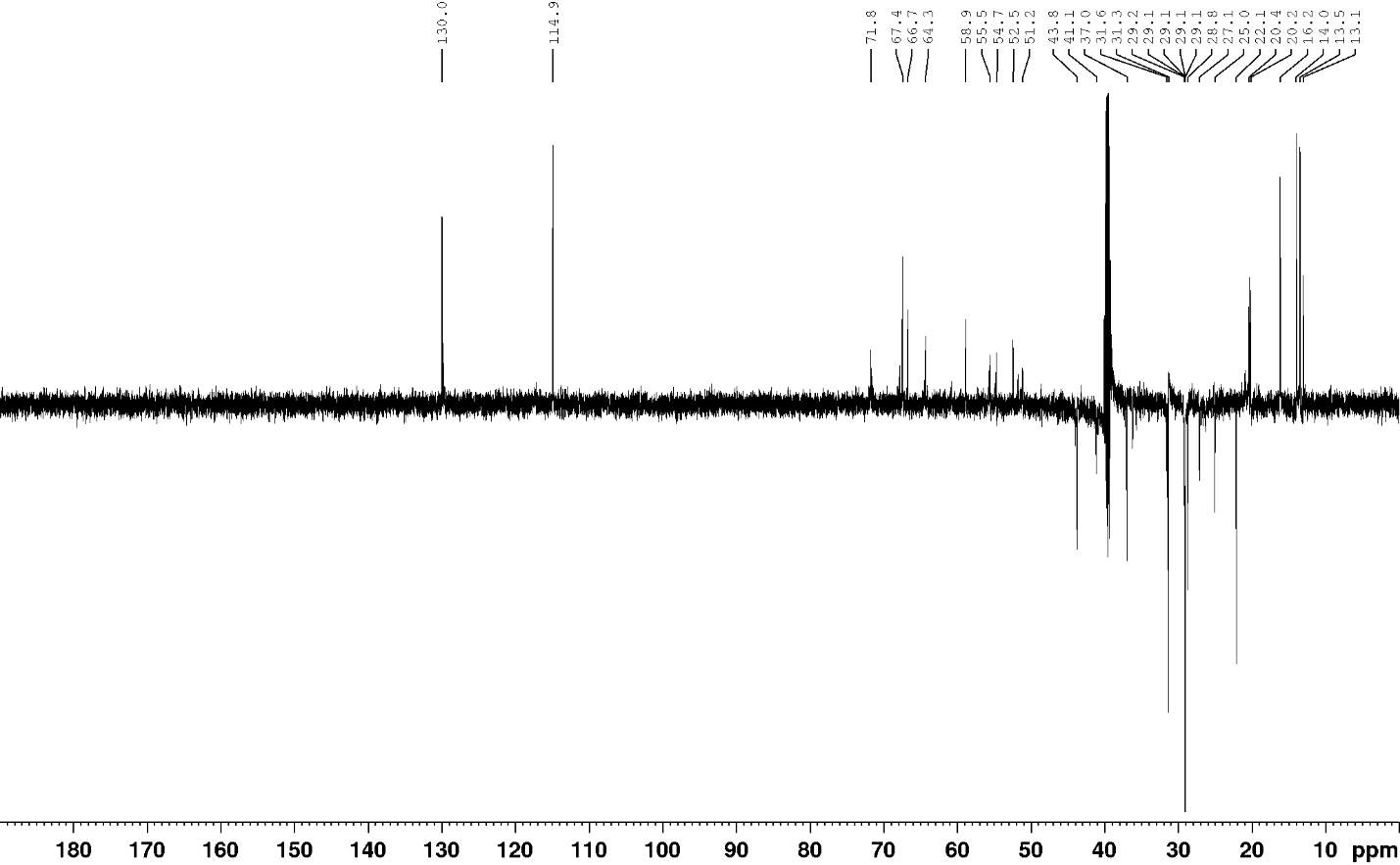

**D**

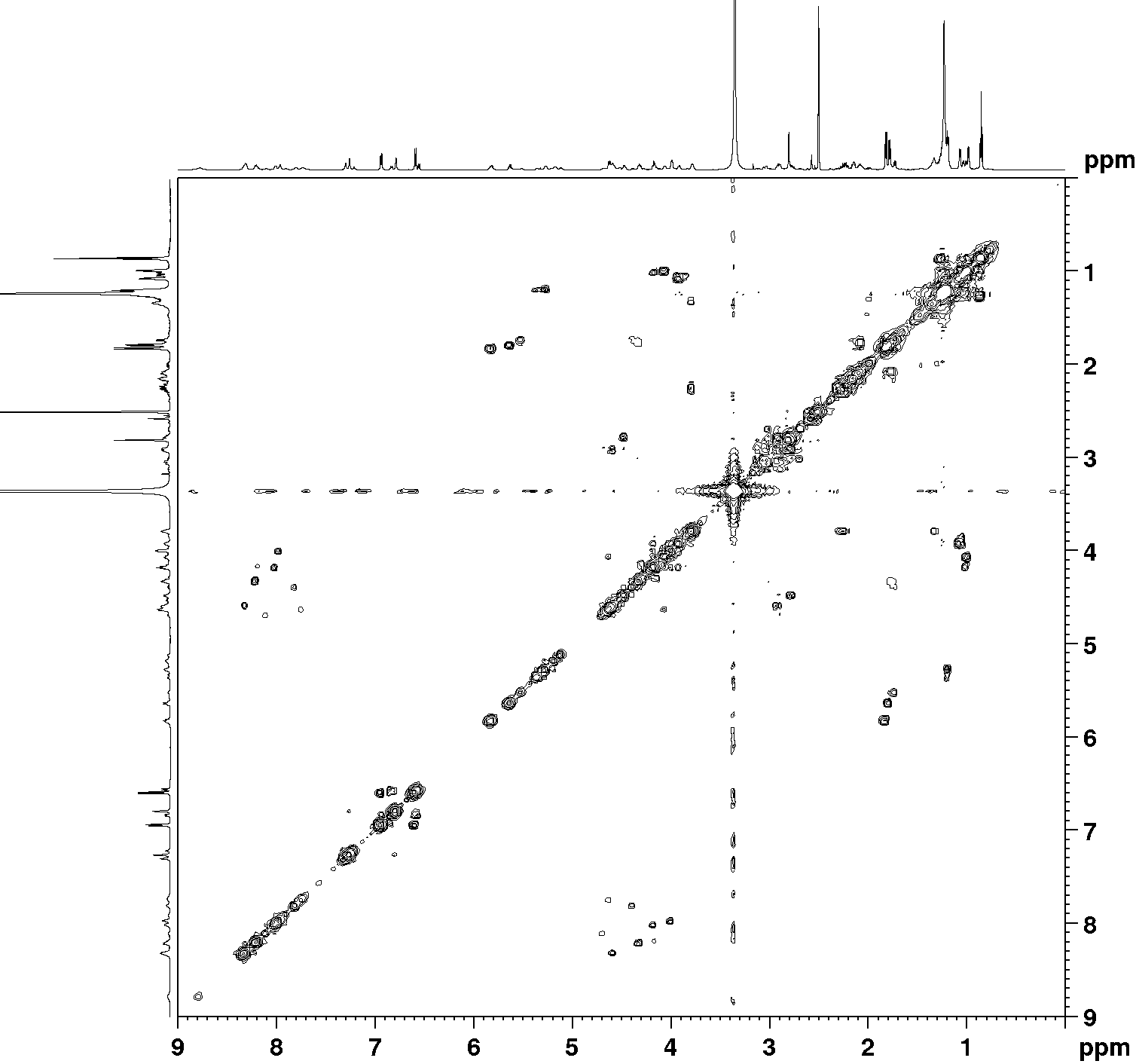

**E**

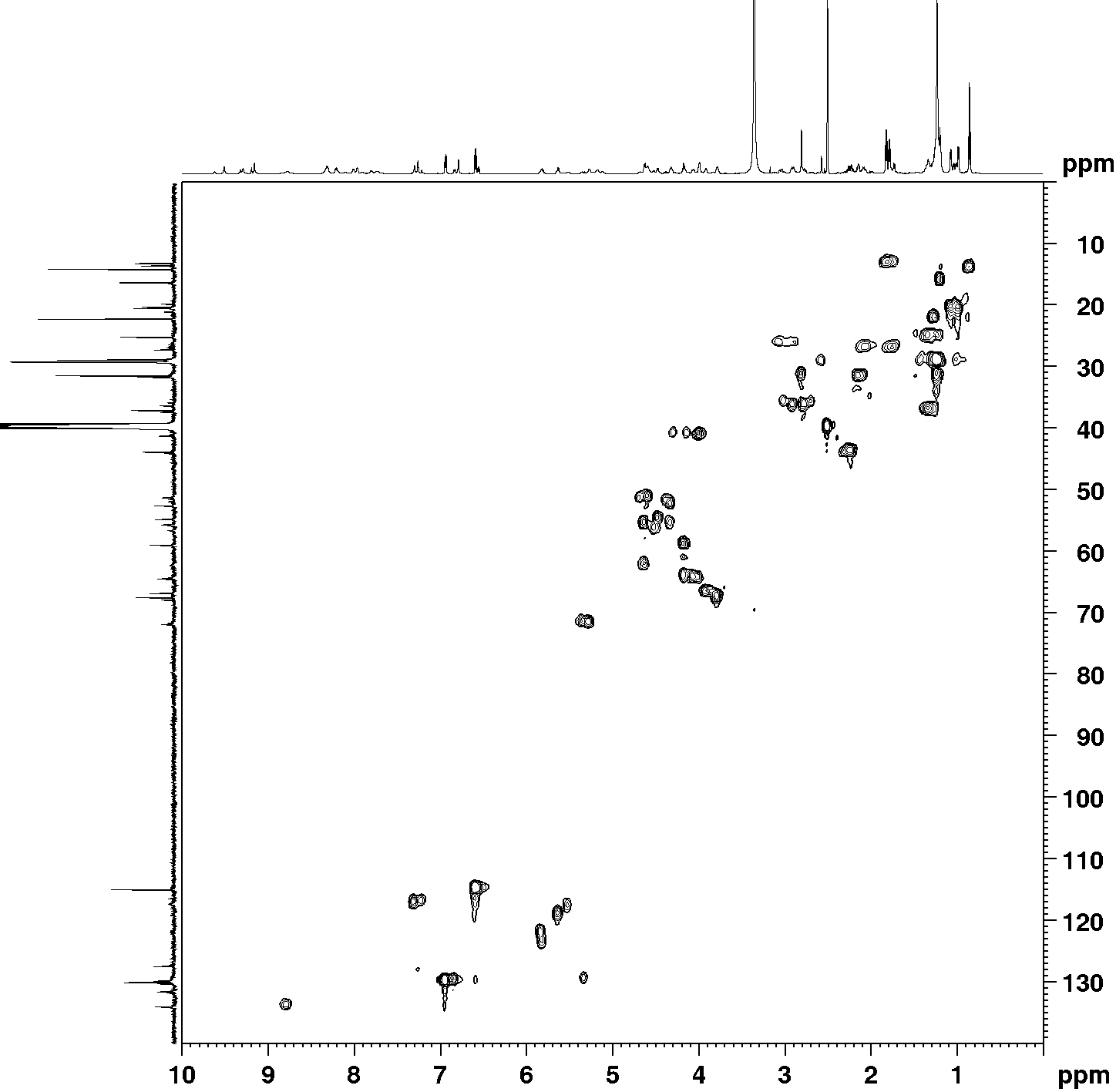

**F**

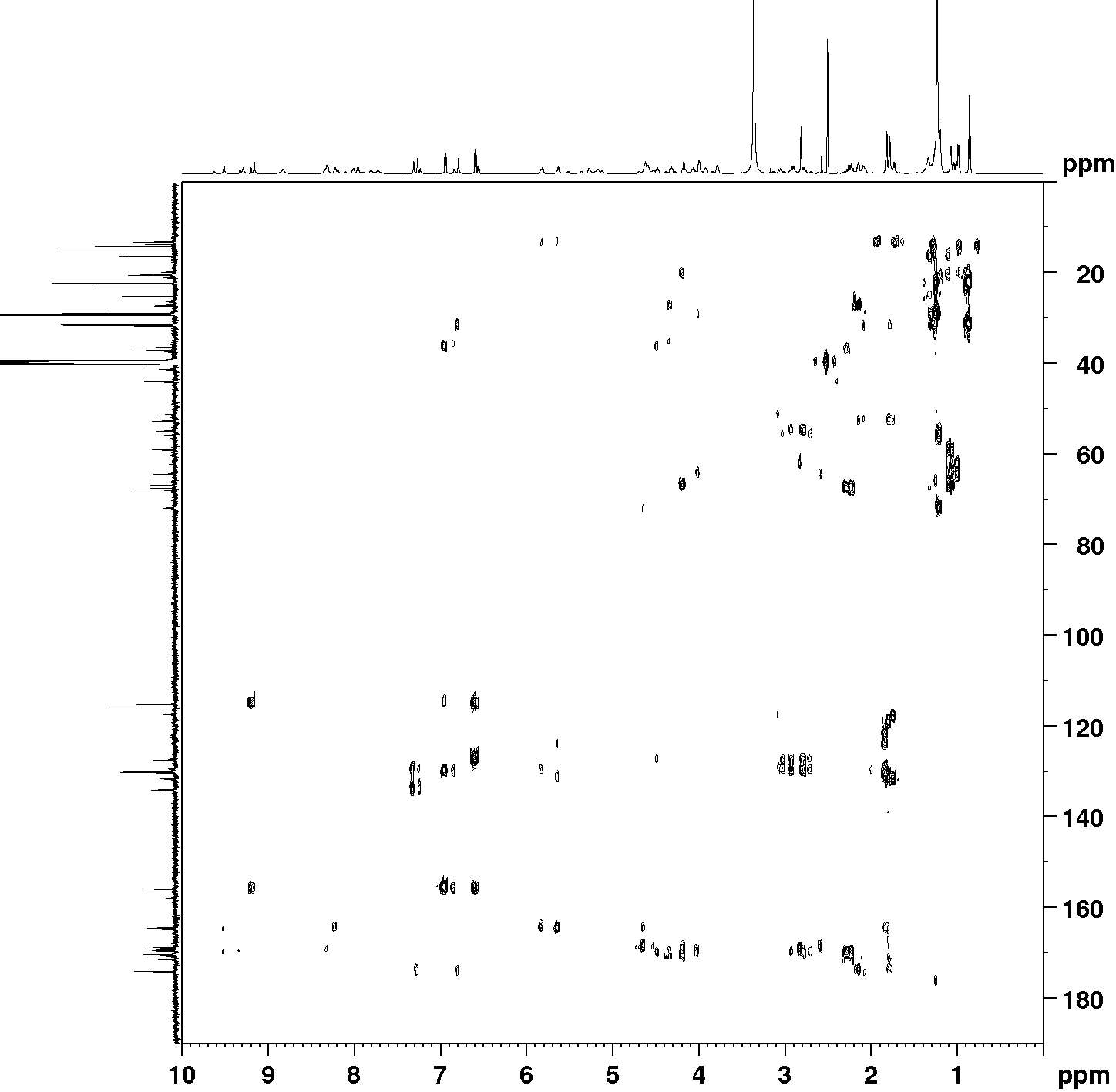

**G**

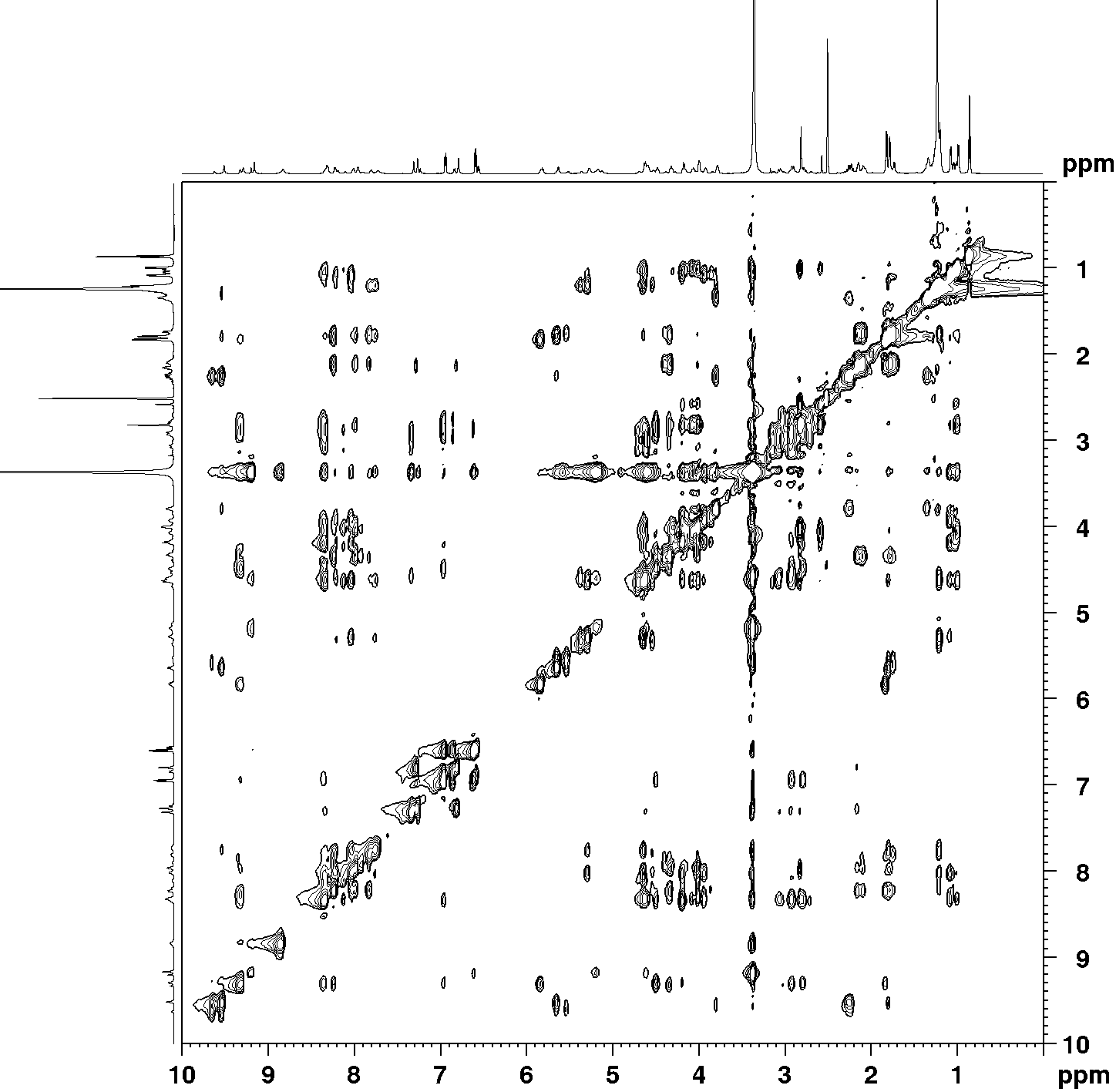

**H**

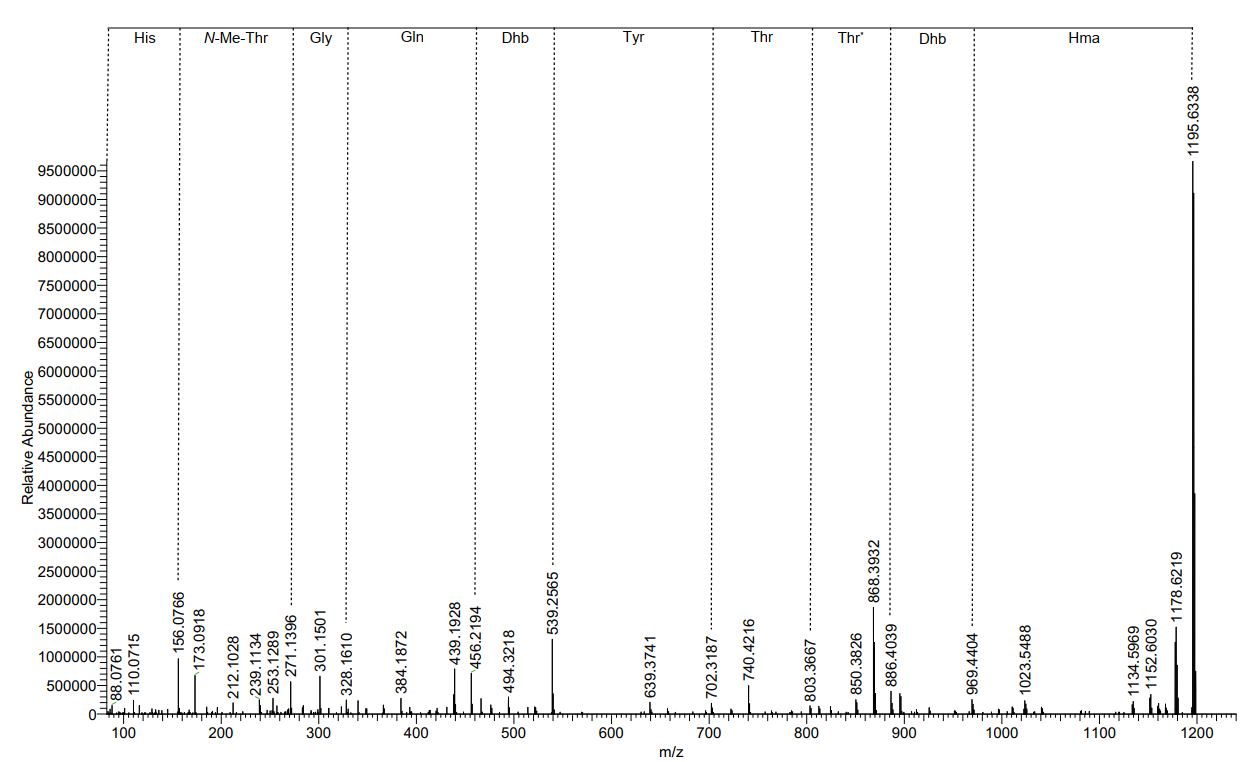

**Figure S4.** Spectra of chromolysin A (**2**). **(A)** ^1^H spectrum. **(B)** ^13^C spectrum. **(C)** DEPT-135 spectrum. **(D)** ^1^H, ^1^H COSY spectrum. **(E)** ^1^H, ^13^C HSQC spectrum. **(F)** ^1^H, ^13^C HMBC spectrum. **(G)** ^1^H, ^13^C NOESY spectrum. **(H)** ESI-MS²-fragmentation. Abbreviations: Dhb, dehydrobutyrine; Hma, β-hydroxymyristic acid; Thr*: mass shift corresponds to loss of C_4_H_5_NO (calc. *m*/*z* 83.0377) due to cleavage of ester bond.

**A**

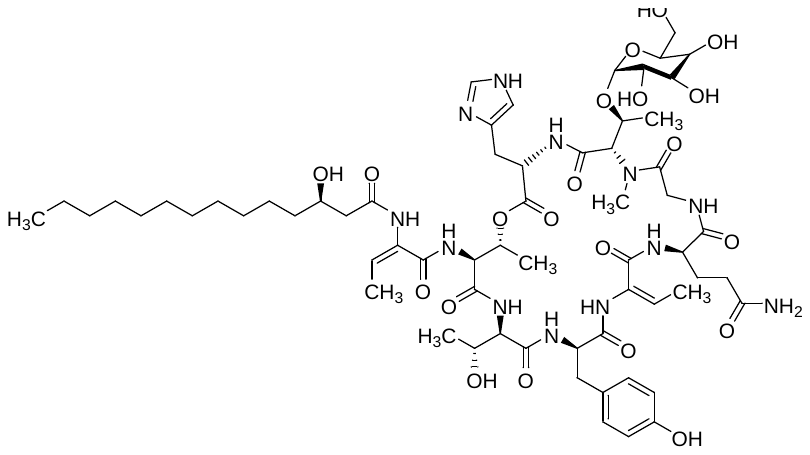

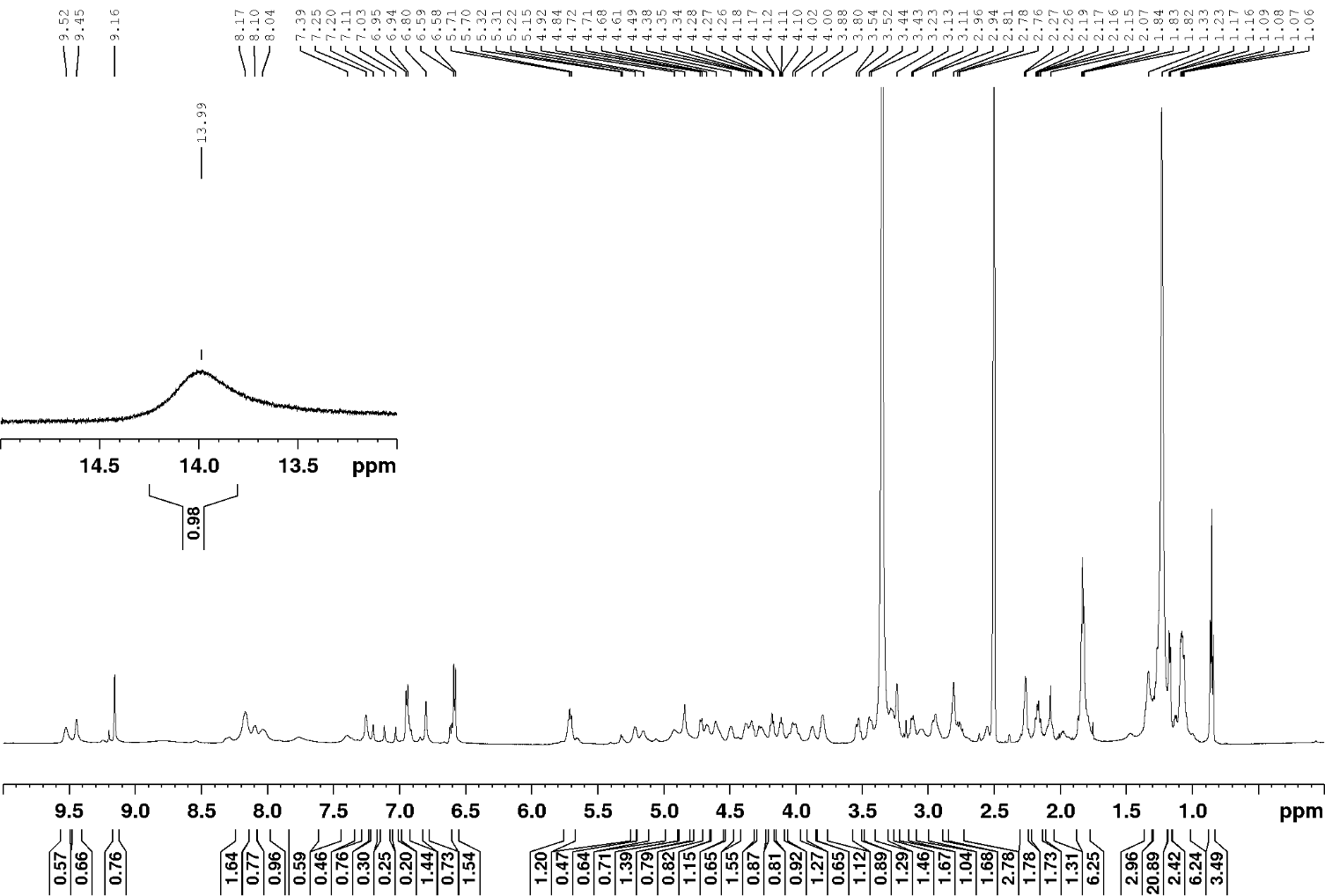

chromolysin B (**3**) (*m*/*z* 1357.6886 [*M*+H]^+^)

**B**

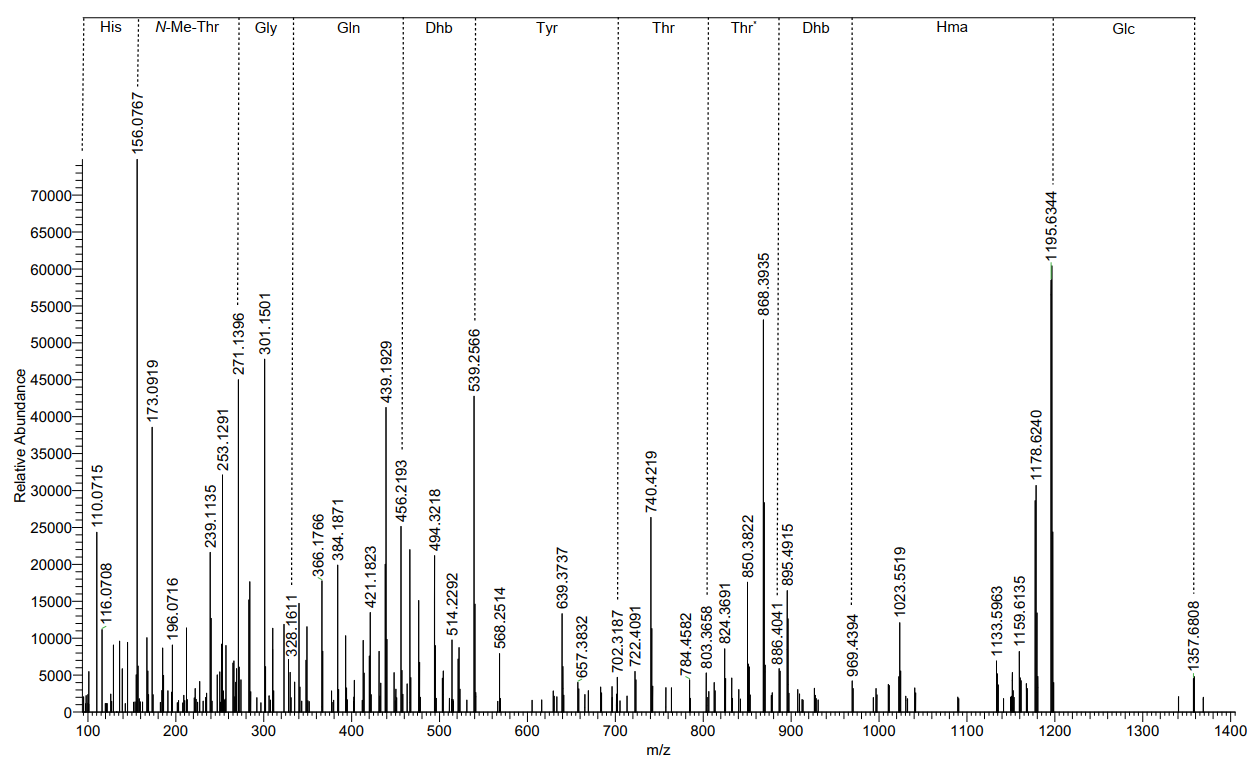

**Figure S5.** Proposed structure of chromolysin B (**3**). **(A)** ^1^H-NMR spectrum. **(B)** ESI-MS²-fragmentation. Abbreviations: Glc, glucose.

**A**

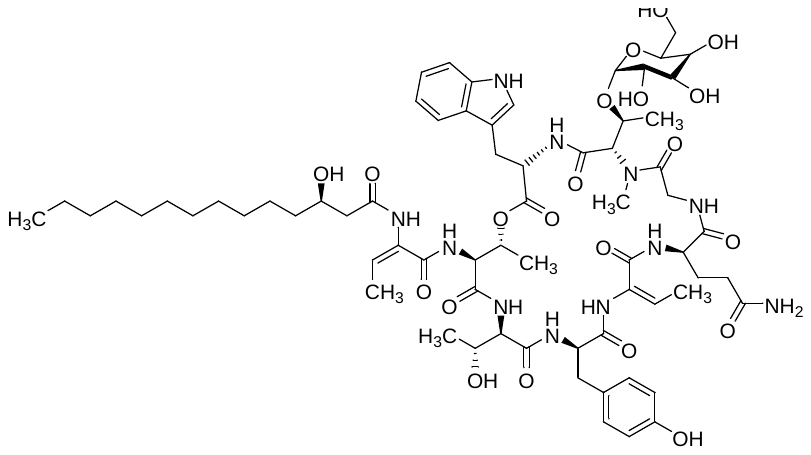

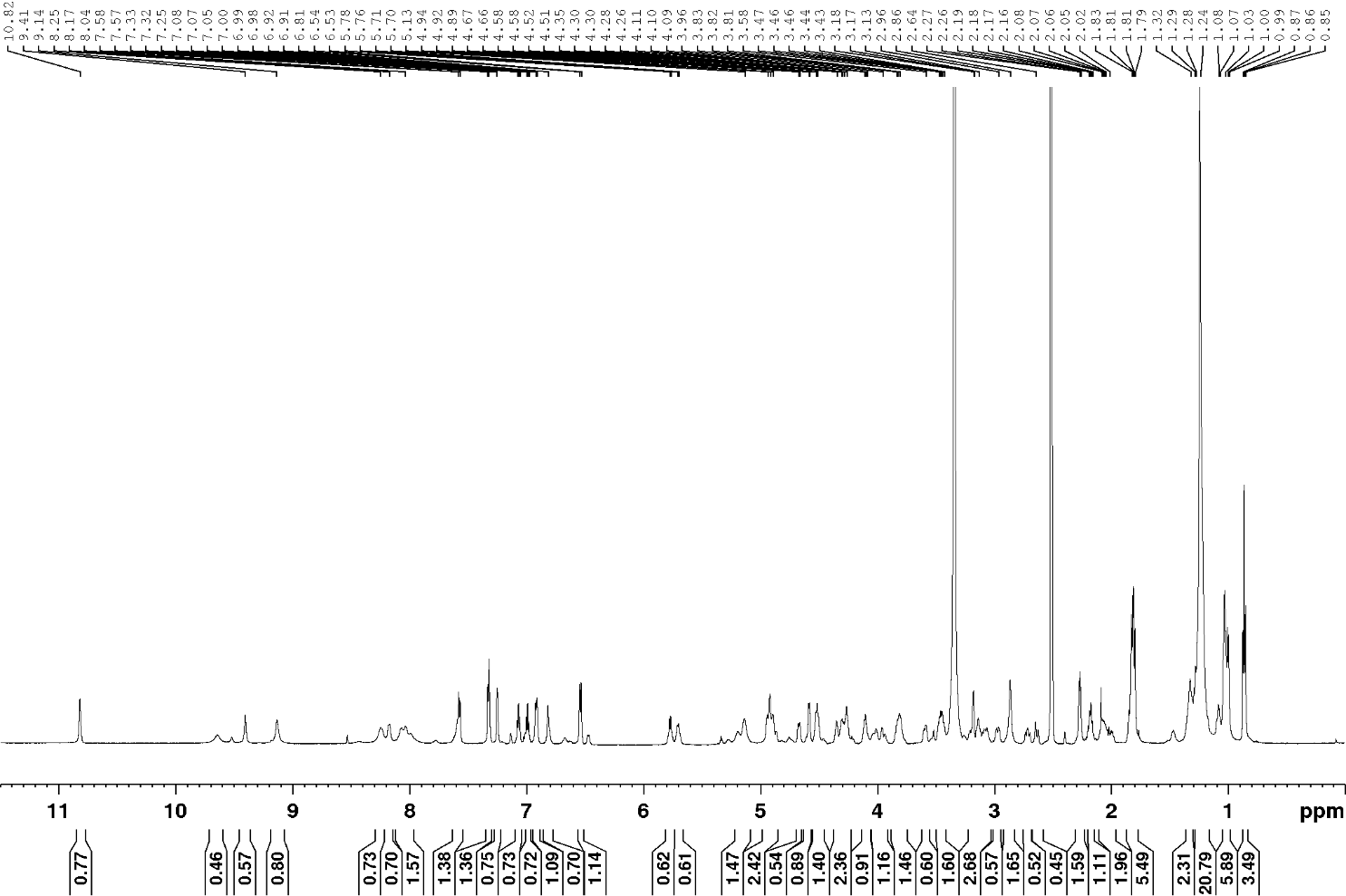

chromolysin C (**4**) (*m*/*z* 1406.7090 [*M*+H]^+^)

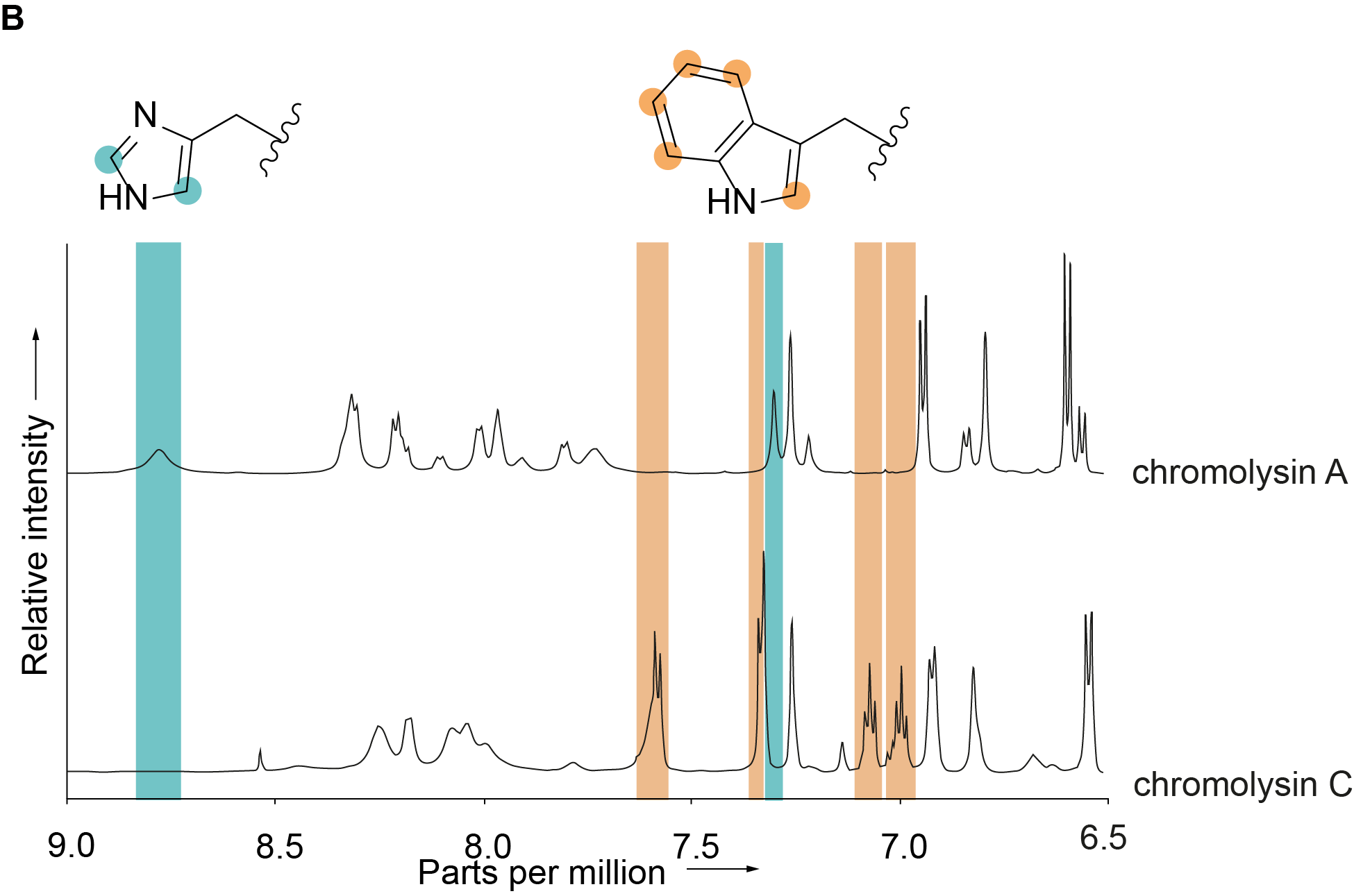
**C**

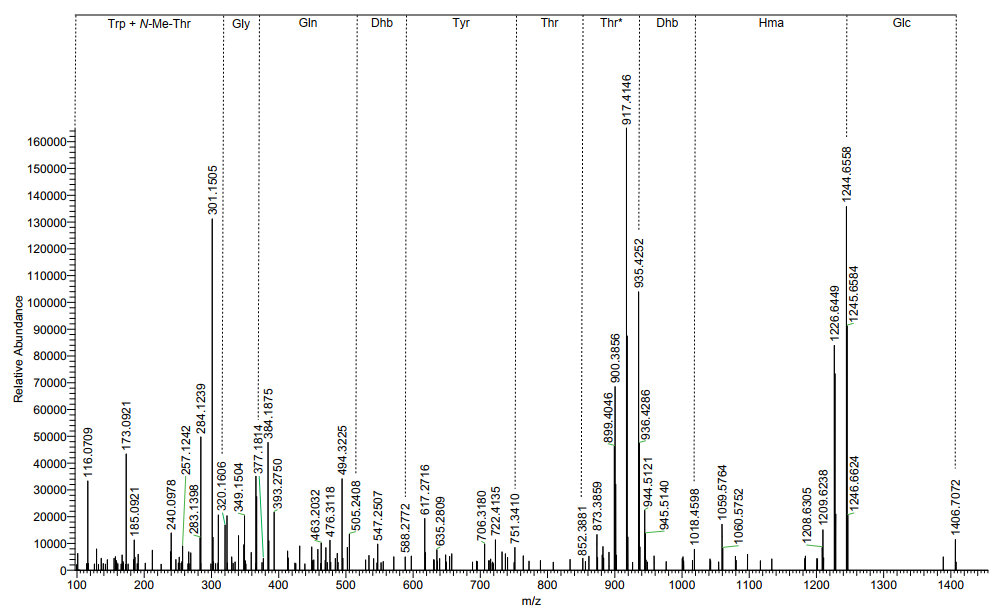
**D**

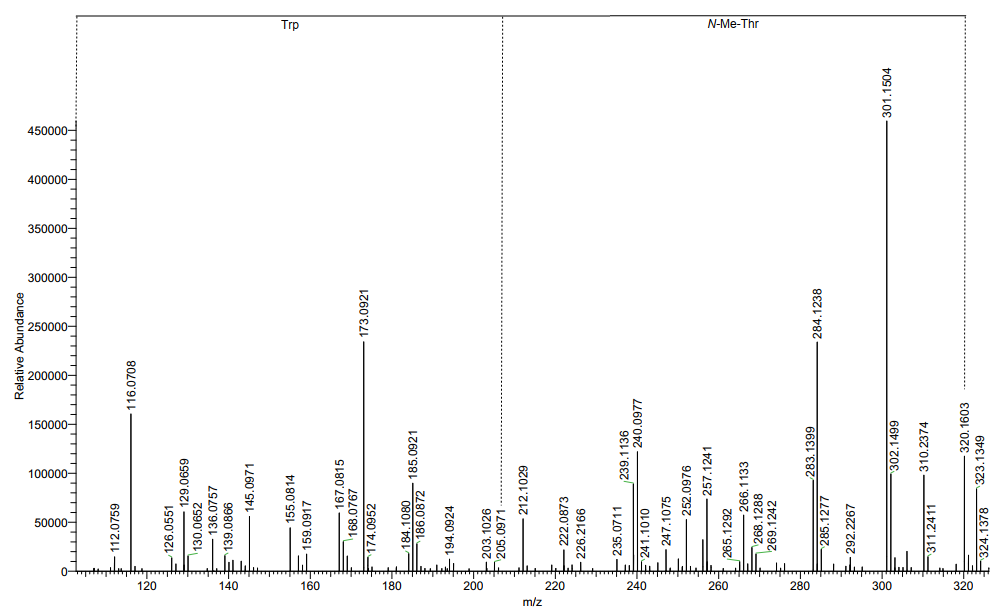

**Figure S6.** Proposed structure of chromolysin C (**4**). **(A)** ^1^H-NMR spectrum. **(B)** Comparison of ^1^H-NMR signals of **2** and **4** in the aromatic region (9.00–6.50 ppm). Coloured peaks correspond to His and Trp, respectively. **(C)** ESI-MS²-fragmentation at 20 eV collision energy. No Trp fragment ion is detected. **(D)** ESI-MS²-fragmentation at 20 eV collision energy. Trp fragment ion is detected. Mass range: *m*/*z* 100.0000–325.0000.

**A**

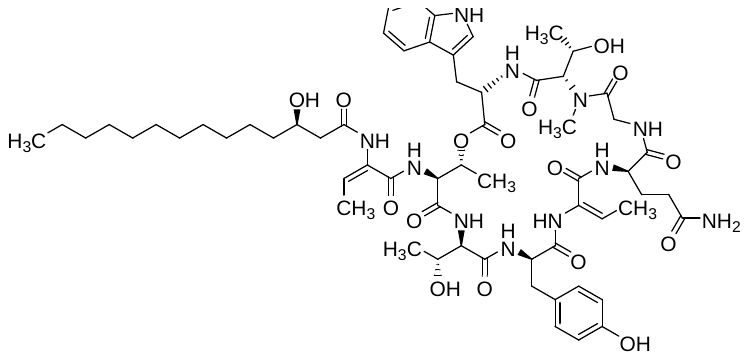

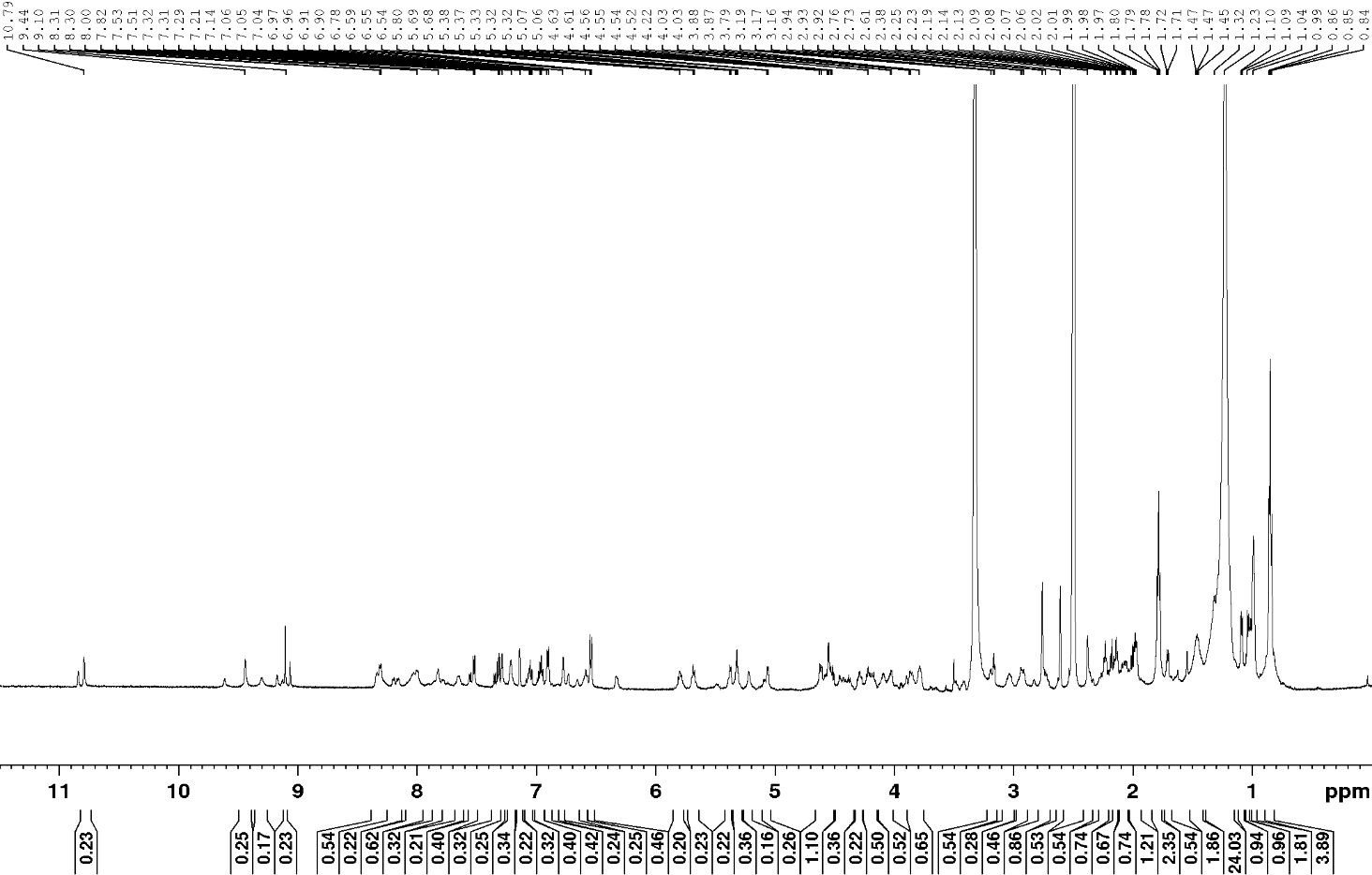

chromolysin D (**5**) (*m*/*z* 1244.6562 [*M*+H]^+^)

**B**

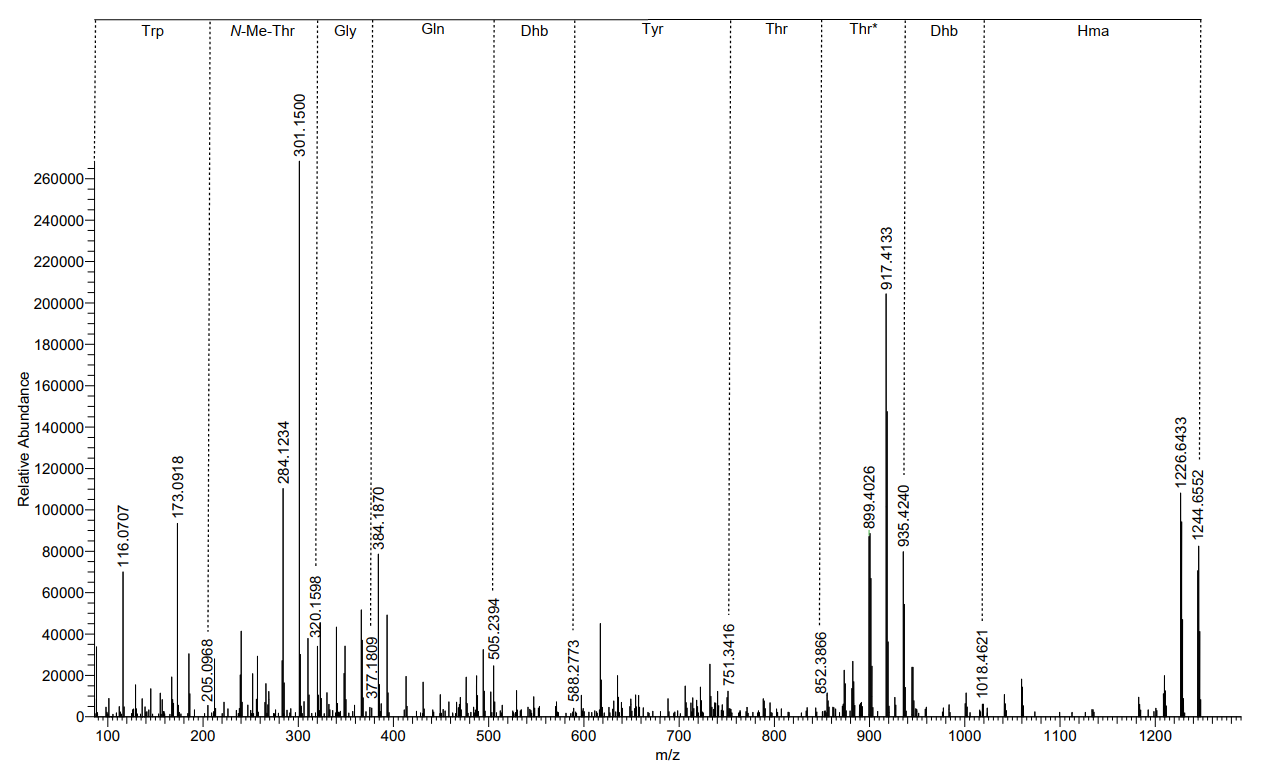

**Fig S7.** Proposed structure of chromolysin D (**5**). **(A)** ^1^H-NMR spectrum. **(B)** ESI-MS²-fragmentation.

**A**

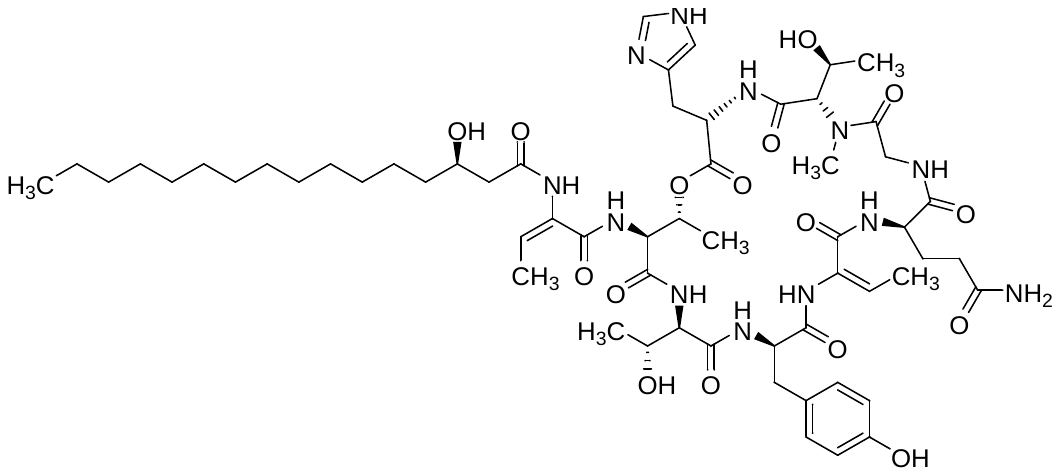

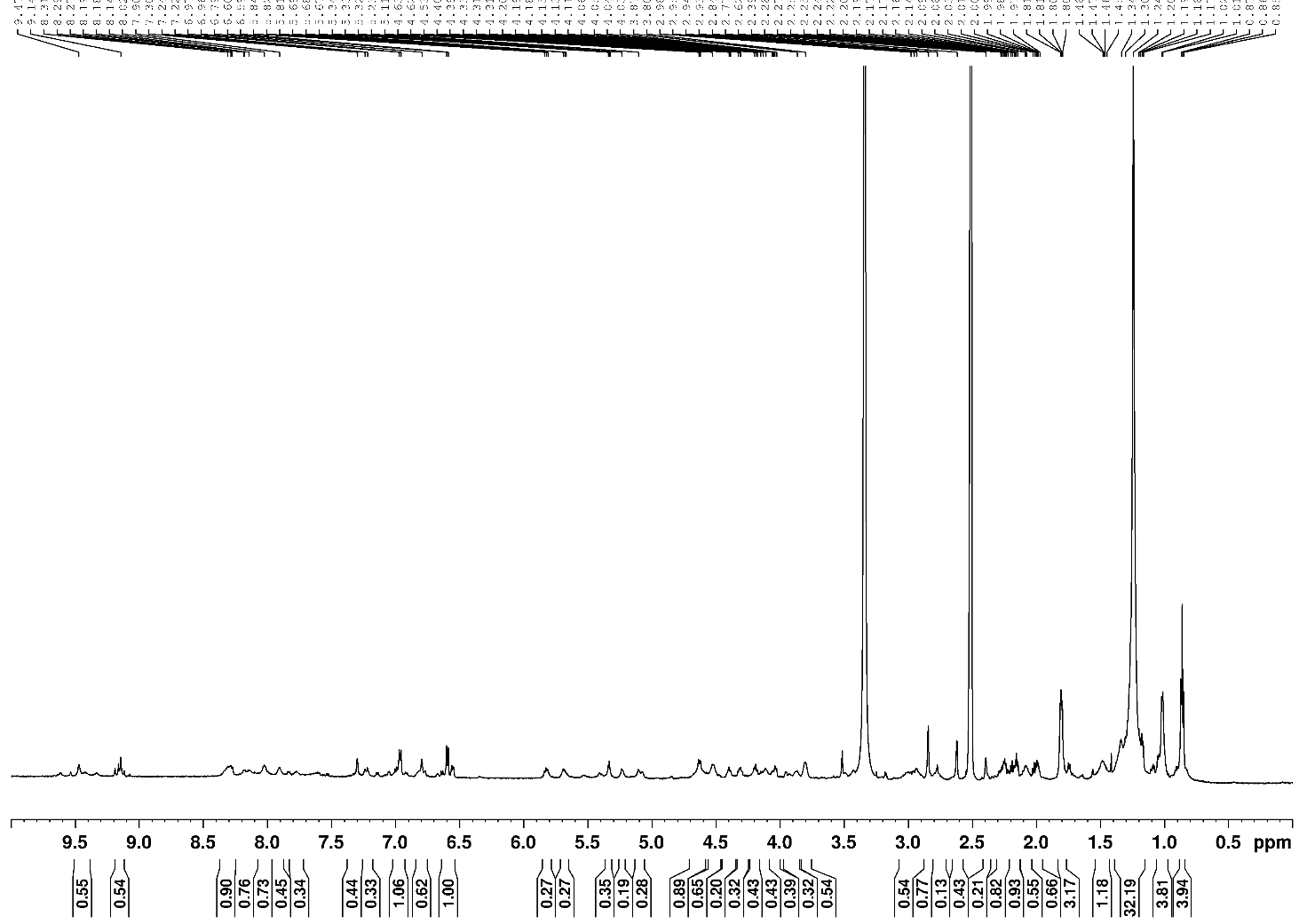

chromolysin E (**6**) (*m*/*z* 1223.6671 [*M*+H]^+^)

**B**

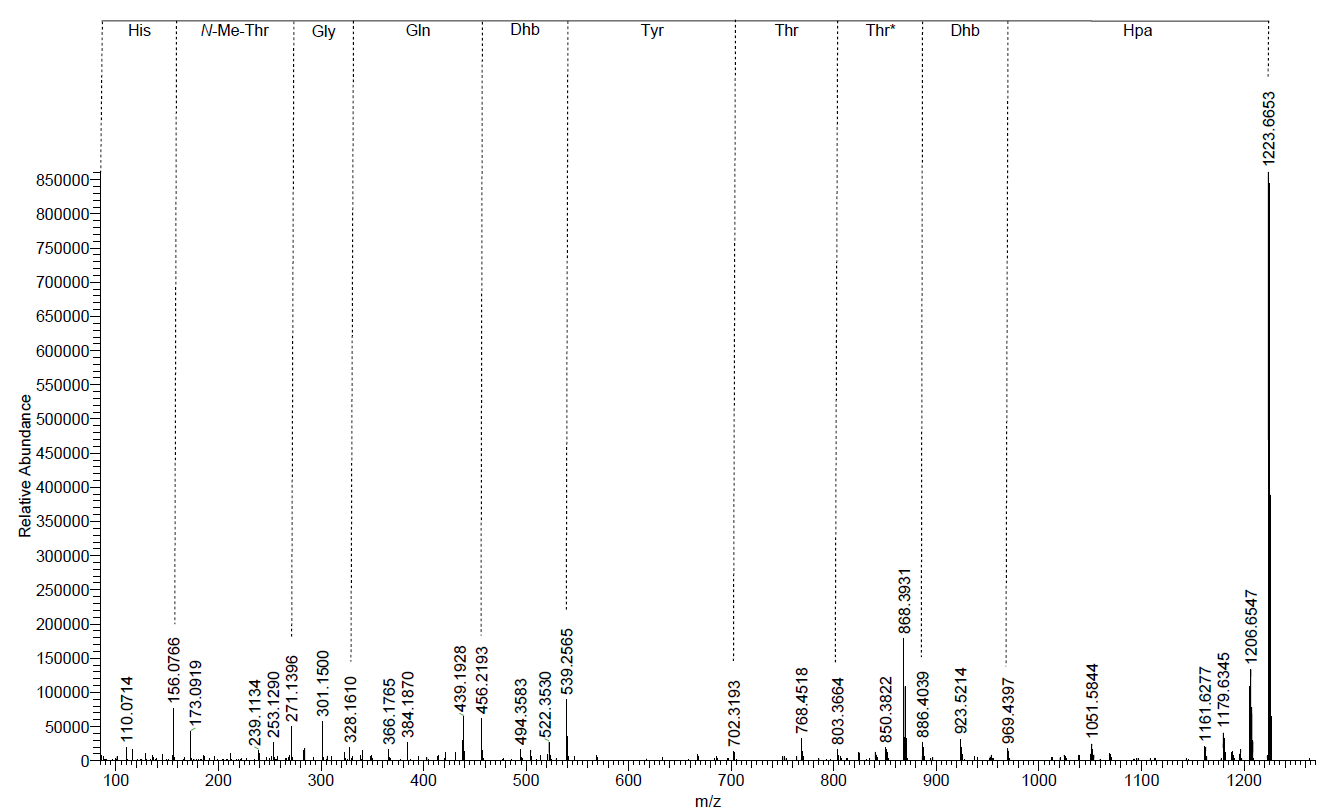

**Fig S8.** Proposed structure of chromolysin E (**6**). **(A)** ^1^H-NMR spectrum. **(B)** ESI-MS²-fragmentation. Abbreviations: Hpa, β-hydroxypalmitinic acid.
